# A circuit from the locus coeruleus to the anterior cingulate cortex modulates offspring interactions in mice

**DOI:** 10.1101/2022.12.04.519053

**Authors:** Alberto Corona, Jane Choe, Rodrigo Muñoz-Castañeda, Pavel Osten, Stephen D. Shea

## Abstract

Social sensitivity to other individuals in distress is crucial for survival. The anterior cingulate cortex (ACC) is a structure involved in making behavioral choices and is influenced by observed pain or distress. Nevertheless, our understanding of the neural circuitry underlying this sensitivity is incomplete. Here, we reveal unexpected sex-dependent activation of ACC when parental mice respond to distressed pups by returning them to the nest (‘pup retrieval’). We observe sex differences in the interactions between excitatory and inhibitory ACC neurons during parental care, and inactivation of ACC excitatory neurons increased pup neglect. Locus coeruleus (LC) releases noradrenaline in ACC during pup retrieval, and inactivation of the LC-ACC pathway disrupts parental care. We conclude that ACC maintains sex-dependent sensitivity to pup distress under LC modulation. We propose that ACC’s involvement in parenting presents an opportunity to identify neural circuits that support sensitivity to the emotional distress of others.

## INTRODUCTION

The ability to detect when others are experiencing distress or danger, and to assist them, is a fundamental aspect of human and animal social behavior. However, ideas about the neural circuit substrates of such behavior and its evolutionary antecedents remain speculative. Parental care is one common scenario that requires an organism to weigh its behavioral options in terms of their risk to itself and the potential benefits to another animal. Parenting offers little proximal benefit to the caregiver, yet protecting offspring when they are in potential peril is essential for the survival of the species ^1^. This decision is often heavily influenced by the sex of the parent. In many species, males and females display dramatic differences in parenting behaviors ^2^. For instance, female mammals typically care for the offspring, whereas males are often aggressive toward the young (reviewed in^1–5)^ with only 5-10% of mammalian species exhibiting paternal care ^6^.

Thanks to recent work on the neural circuits involved in pro-social behaviors in rodents, we are building a better understanding of the potential neural basis of social sensitivity to distress ^7–12^. For example, the anterior cingulate cortex (ACC) responds to social information and the emotional state of others, especially distress ^7, 10^. ACC has been implicated in social transfer of pain ^10^, cost-benefit decision-making ^13^, and observational fear learning ^7^. Hypoactivity of ACC pyramidal neurons appears to contribute to disrupted social interactions in the *Shank3* mouse model of autism ^14^. With regard to parental behavior, ACC lesions in rats disrupt maternal behavior in early postnatal days (PNDs) ^15^, and fMRI studies in humans have shown that ACC is activated by infant cries ^16^. Taken together, these data suggest that ACC is an important regulator of social cognition and may participate in computations that balance the drive to help others against the drive to avoid danger and risk.

One potentially relevant input to ACC is locus coeruleus (LC), which consists of a bilateral pair of nuclei in the pons that serve as nearly the sole source of noradrenaline (NA) for the brain. LC is an important regulator of stress, arousal, and state-dependent cognitive processes ^17, 18^, and it also has an established role in maternal behavior ^19^. Mice lacking a gene necessary for NA synthesis exhibit severe deficits in maternal behavior, with most of the pups dying due to maternal neglect. This deficit is obviated by restoring NA signaling just before the birth of the pups ^19^.

When mouse pups are separated from the nest, they are at risk and emit ultrasonic distress vocalizations (USVs; 50 – 80kHz). In response, the dam returns them to the nest in a behavior called ‘pup retrieval’ ^20–25^. Recent work from our laboratory shows that a large fraction of LC neurons becomes robustly and precisely active as dams retrieve wayward pups and return them to the nest ^26^. We proposed that LC contributes to goal-directed action selection during parental behavior with widespread release of NA ^26^. However, the downstream targets through which LC acts to modulate pup retrieval are unknown. ACC receives robust projections from LC ^27–29^, and the activity in the two regions is highly correlated when external stimuli trigger phasic activation of LC ^30^. Therefore, we hypothesized that LC influences ACC to regulate maternal care.

Although pup retrieval behavior has been observed in sires ^31–34^, their behavior is less robust and consistent compared to dams. In particular, sires are slower at retrieving pups to the nest. We hypothesize that sires are less sensitive to offspring distress compared to dams. To investigate potential sex differences in neural circuits that control sensitivity to offspring distress, here we compared how dams and sires respond to distressed pups that are outside of the nest. We used brain-wide imaging of the immediate early gene *c-fos* to compare patterns of brain activity between retrieving and non-retrieving dams and sires. These experiments uncovered previously unappreciated sex-dependent activation of ACC during interactions with pups. Fiber photometry recordings from parents actively engaged in pup retrieval corroborated this finding, revealing that excitatory and inhibitory neurons in ACC show stronger reciprocal activity patterns during pup retrieval behavior in dams as compared to sires. Furthermore, chemogenetic inactivation of excitatory neurons in ACC increases the latency to retrieve pups and decreases parental responsiveness to interact with pups in distress. We confirm the existence of a projection from LC to ACC, and we found that phasic firing events in LC evoked by pup retrieval behavior trigger NA release in ACC. Finally, we showed that inactivating ACC inputs from LC increased parental neglect. Therefore, we propose that ACC adjusts parental sensitivity to pup distress through LC modulation in a sex-dependent manner.

## RESULTS

### Sex-dependent differences in pup retrieval behavior

As a first step in our investigation into sex differences in parental behavior and the underlying neural circuits, we observed parental interactions of CBA/CaJ dams and sires with their pups. While both sexes participated in parental care, we observed many quantitative sex differences in the behavior. We quantified the efficacy of pup retrieval in the home cage (Figure 1A) as described ^35^. Briefly, pups were scattered around the home cage, and we calculated a normalized measure (ranging from 0 – 1) of the latency to return all pups to the nest (see Methods). Higher latency is considered poorer performance. Typically, dams exhibited reliable retrieval on early trials and rapidly improved over time (Figure 1B and Figure S1). In contrast, sires were frequently unreliable and inconsistent in retrieval, and they failed to improve as rapidly as dams. Consequently, they had a higher mean latency to gather pups (Figure 1B-E). Previous work showed that sires of the ICR strain do not gather pups in a novel environment ^31^. Therefore, we performed the retrieval assay in a novel cage to assess sex differences in contextual regulation of parental behavior. Consistent with previous reports ^31^, we found that among sires, overall mean latencies and improvement across days were poorer in the novel cage, but this was not the case in dams (Figure 1C).

**Figure 1.**
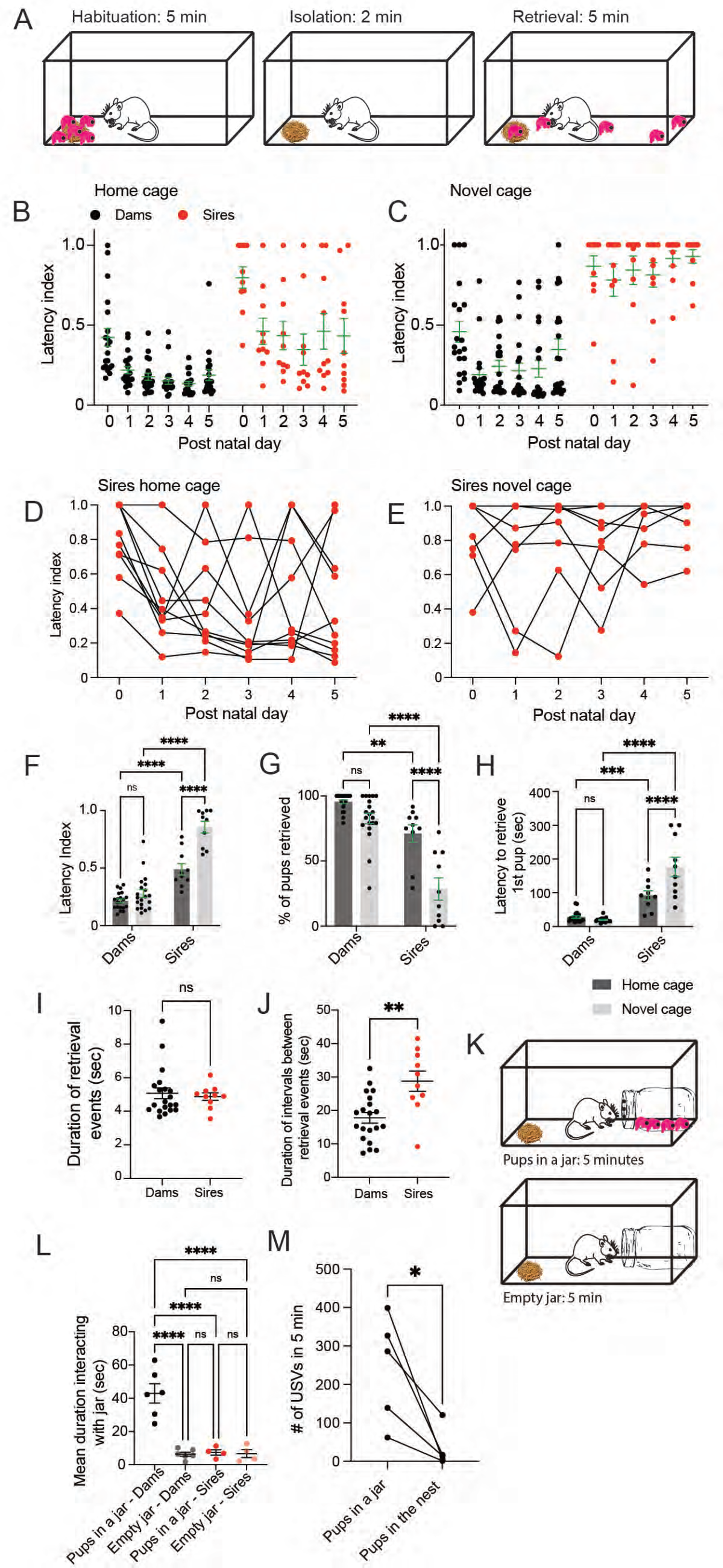
Sex-dependent differences in pup retrieval behavior. **A)** Schematic of behavioral paradigm. **B-C)** Scatterplots showing a normalized measure of latency to gather pups for dams (n = 20 per context), and sires (n = 10 per context) in their home cage or a novel cage, respectively, at postnatal days (PNDs) 0 – 5. Green lines represent mean ± s.e.m. **D)** Plot of retrieval latency of sires in the home cage across days. Lines track each individual’s performance across PNDs 0 – 5. **E)** Plot of retrieval latency of dams in a novel cage across days. Lines track each individual’s performance across PNDs 0 – 5. **F)** Mean latency index for all subjects at PND 0 – 5 for dams and sires. Whiskers represent mean ± s.e.m. (n = 10 sires and 20 dams). A two-way ANOVA was performed on latency to retrieve pups by sex and context. There was a statistically significant interaction between the effects of sex and context on pup retrieval [F(1, 56)=17.28, p = 0.0001]. Holm-Sidak posthoc tests were subsequently performed for individual group comparisons ****p<0.0001. **G)** Plot of percentage of pups retrieved averaged across PNDs 0 – 5 in the home cage and a novel cage. Whiskers represent mean ± s.e.m. (n = 10 sires and 20 dams). A two-way ANOVA was performed on the percentage of pups retrieved by sex and context. There was a statistically significant interaction between the effects of sex and context on percentage of pups retrieved [F(1, 56)=9.592, p = 0.0031]. Holm-Sidak’s post hoc tests were subsequently performed for individual group comparisons. ****p<0.0001; **p=0.0023. **H)** Plot of mean latency to retrieve the first pup at PNDs 0 – 5. Whiskers represent mean ± s.e.m. A two-way ANOVA was performed on the latency to retrieve the first pup by sex and context. There was a statistically significant interaction between the effects of sex and context on latency to retrieve the first pup [F(1, 56)=17.38, p = 0.0001]. Holm-Sidak’s post hoc tests were subsequently performed for individual group comparisons. ****p<0.0001; ***p=0.0003. **I)** Plot of mean duration of retrieval events as measured from the time the subject makes pup contact to the time it drops the nest in the nest in the home cage. (n = 20 dams and n=10 sires, unpaired t test, p=0.6903). **J)** Plot of mean duration of intervals between retrieval events as measured from the end of the previous retrieval event to the start of the next one in the home cage. (n = 20 dams and n=10 sires, unpaired t test, **p=0.0014). **K)** Schematic of jar behavioral paradigm. **L)** Plot of mean duration of time that subjects spent interacting with pups in a jar versus an empty jar. Green lines represent mean ± s.e.m. An ordinary one-way ANOVA was performed [F (25.41) p<0.0001], followed by Tukey’s post hoc tests for individual group comparisons. ****p<0.0001. **M)** Plot of the number of USVs emitted by the pups during a 5-minute session in the jar compared with a 5-minute session in the nest without the dam. Paired t-test *p=0.0262.

Across post-natal days (PNDs) 0 – 5, sires retrieved fewer pups than dams in both contexts (Figure 1G), and sires took longer than dams to initiate contact with the first pup in both contexts (Figure 1H). The duration of individual retrieval events, measured as the time from pup contact to the time the pup was deposited in the nest, did not differ between sires and dams (Figure 1I), demonstrating that sires and dams are equally capable of performing motor aspects of the behavior. However, the intervals between retrieval events were significantly longer in sires compared to dams (Figure 1J). Regardless of the context, pup retrieval performance was always significantly poorer in sires as compared to dams (Figure 1F).

We found that dams and sires display differential sensitivity to pups in distress by measuring the time dams and sires spent trying to interact with inaccessible isolated pups. We trapped the pups in a glass jar and placed it in the home cage. The jar had a plastic lid with small holes on it, so the animals were able to hear and smell the pups, but they were not able to touch them (Figure 1K). Being trapped in the jar elicited overt signs of distress in the pups; the pups emitted significantly more USVs when they were trapped compared to when they were in the nest (Figure 1M). We recorded each animal’s behavior in their home cage for 5 minutes with the pups trapped in the jar and for 5 minutes with an empty jar and the pups absent. Dams spent considerable time interacting with the sealed opening of the jar, including sniffing, biting, and clawing at the lid (Supplementary Movie 1). Trapping the pups in the jar elicited a significantly greater investigatory response in the dams when compared to an empty jar or males with either jar (Figure 1L).

### Brain-wide activity mapping reveals regions associated with pup retrieval

Having confirmed the expected sex differences in parenting behavior, we asked whether there are corresponding sex differences in the brain-wide patterns of neural activation when parents interact with their pups in distress. To identify regions that modulate parental interactions in dams and sires, we performed a screen for differential expression of the immediate early gene *c-fos* with an automated pipeline for brain-wide *c-fos* mapping ^36^. In short, *c-fos* expression was induced by one of several different behavioral conditions in dams and sires, and then subjects were sacrificed after 90 minutes and perfused. Brains were cleared, immunolabeled, and imaged with light-sheet microscopy, and *c-fos*^+^ nuclei were automatically counted with custom software (Figure 2A).

**Figure 2.**
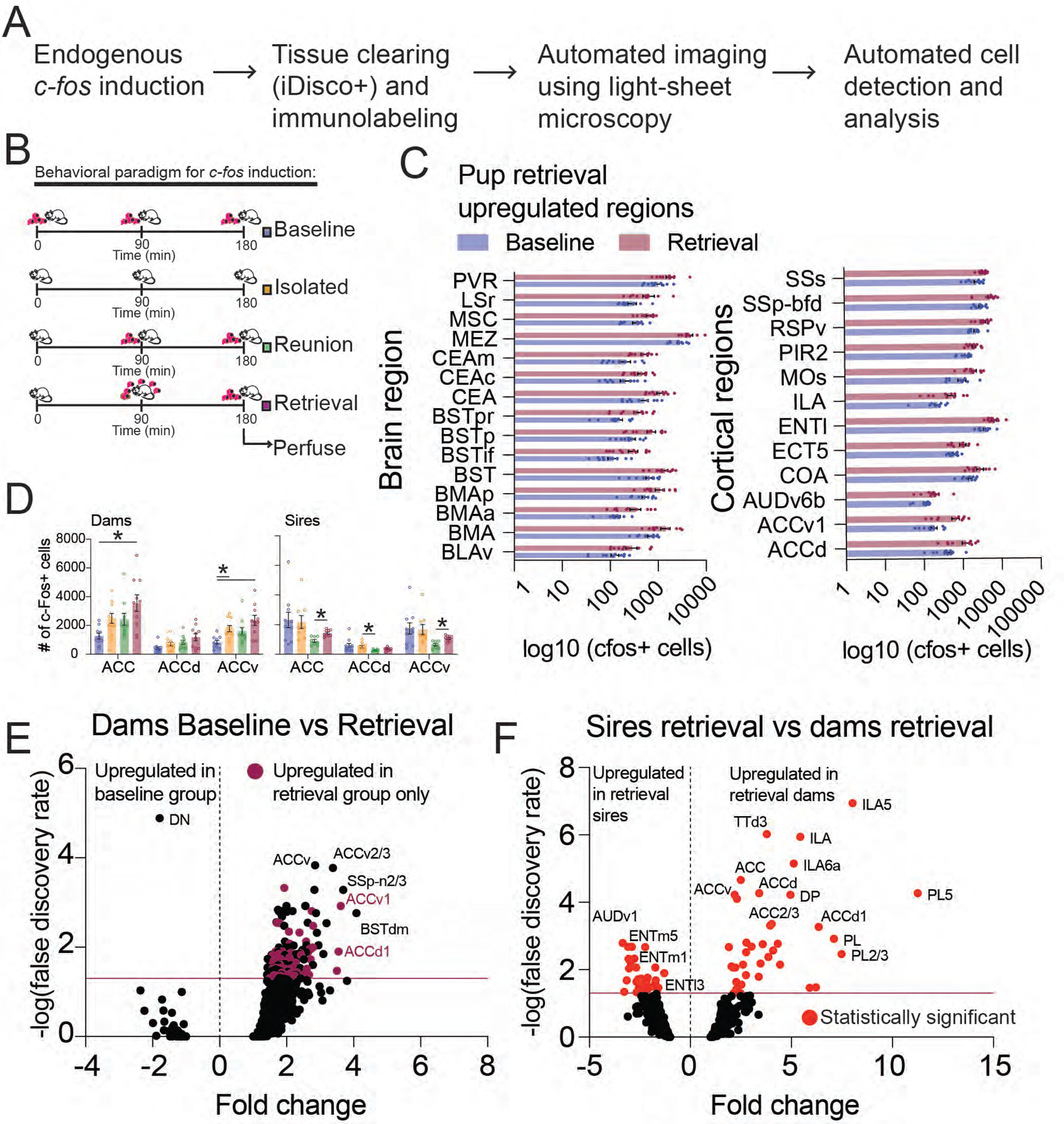
Brain-wide *c-fos* expression screen. **A)** Brain-wide *c-fos* expression mapping workflow. **B)** Schematic representation of behavioral paradigm to induce *c-fos* expression. There were four groups in the assay: undisturbed, isolated, reunion, and retrieval. The experiment was performed in dams and sires. All experiments were performed in the dark during the light cycle in a soundproof behavioral box. All mice were perfused immediately after the completion of the experiment. **C)** Plots of brain regions in which *c-fos* expression was uniquely upregulated in the retrieval vs baseline comparison in dams. PVR: Periventricular region; LSr: Lateral septal nucleus, rostral part; MSC: Medial septal complex; MEZ: Medial hypothalamic zone; CEAm: Central amygdala, medial part; CEAc: Central amygdala, capsular part; CEA: Central amygdala; BSTpr: Bed nucleus of the stria terminalis, posterior division, principal nucleus; BSTp: Bed nucleus of the stria terminalis, posterior division; BSTif: Bed nucleus of the stria terminalis, posterior division, interfascicular nucleus; BST: Bed nucleus of the stria terminalis; BMAp: Basomedial amygdalar nucleus, posterior part; BMAa: Basomedial amygdalar nucleus, anterior part; BMA: Basomedial amydalar nucleus; BLAv: Basolateral amygdalar nucleus, ventral part; SSs: Supplemental somatosensory area; SSp-bfd: Primary somatosensory area, barrel field; RSPv: Retrosplenial area, ventral part; PIR2: Piriform area, pyramidal layer; MOs: Secondary motor area; ILA: Infralimbic area; ENTl: Entorhinal area, lateral part; ECT5: Ectorhinal area/Layer 5; COA: Cortical amygdalar area; AUDv6b: ventral auditory area, layer 6b; ACAv1: Anterior cingulate area, ventral part, layer 1; ACAd: Anterior cingulate area, dorsal part. **D)** Plot of *c-fos*+ cells in the anterior cingulate cortex of dams and sires in response to pup interactions (dams n = 10 per group; sires n = 10 baseline, 9 isolated, 8 reunion, and 9 retrieval Tukey’s multiple comparison test *p<0.05) **E)** Volcano plot of *c-fos* induction in regions of the baseline vs retrieval conditions in dams as fold change in *c-fos*+ cells. False discovery rate (FDR) analysis was carried out by the Benjamini-Hochberg procedure. Purple data points correspond to statistically significant ROIs unique to comparing baseline vs retrieval groups and no other control groups. The purple horizontal line indicates the significant threshold with an FDR of 0.05. All ROIs above the line are statistically significant. **F)** Volcano plot showing the statistical comparisons of retrieval groups between dams and sires. All positive fold changes indicate an upregulation in dams and negative fold changes indicate an upregulation in sires. Significant ROIs are depicted in red. The horizontal purple line depicts the significant threshold with an FDR of 0.05.

We examined whole-brain *c-fos* expression patterns in mice that were exposed to one of four conditions: baseline, isolated, reunited with their pups, and pup retrieval (Figure 2B). In dams, considering total *c-fos*^+^ cell counts among all regions, the baseline group showed significantly lower counts and less variability between individuals when compared to the isolated, reunion, and retrieval groups (Figure S2B). Sires showed greater overall variability in *c-fos*^+^ cell counts, including baseline, and no significant differences across most of the groups (Figure S2C and S4E-H). Comparison of *c-fos* expression between the four experimental groups identified brain areas affected by the different behavioral conditions (Figure S3). Therefore, the *c-fos* expression patterns observed in this study result from the presence of the pups, the absence of the pups, the reunion with the pups, or the retrieval of the pups to the nest.

The comparison between dams of the baseline group versus the retrieval group likely reflects changes in regions associated with pup retrieval. We identified brain regions that showed significantly greater or lesser *c-fos* expression in retrieval mice compared to all other control conditions and corrected for false discovery rate (Figure 2C). We observed more *c-fos*^+^ cells in retrieval mice in brain areas that have been previously implicated in parenting. For example, the bed nucleus of the stria terminalis (BNST) ^37–41^, the medial hypothalamic zone, which includes the medial preoptic area, (MEZ) ^32, 33, 42–46^, the medial septal complex (MSC) ^47^, the anterior cingulate cortex (ACC) ^15^, the basomedial amygdala (BMA) ^48, 49^, and the central amygdala (CEA) ^50^. In contrast to the sires (Figure S4), most changes observed in the comparison between the baseline and the retrieval groups in the dams showed an increase in *c-fos*^+^ cells in mice from the retrieval group (Figure 2E), suggesting that the changes in *c-fos* were driven by pup retrieval. We then compared dams and sires from the retrieval groups and identified brain areas with significantly more *c-fos* in dams or sires. Brain areas upregulated in dams relative to sires were primarily regions in the prefrontal cortex including the infralimbic, prelimbic, and cingulate cortices (Figure 2F).

One region that stood out was ACC, which exhibited some of the highest *c-fos* upregulation in dams from baseline to retrieval (Figure 2E) and was upregulated in sires comparing the retrieval condition to the reunion group (Figure 2D). Although ACC lesions impair maternal behavior in rats ^15^, ACC is not widely appreciated as a major regulator of maternal behavior, and the specific function and timing of its involvement in pup retrieval is unexplored. Interestingly, ACC was one of the most upregulated regions in dams relative to sires, potentially indicating sex-dependent modulation of ACC during interactions with pups (Figure 2F). We chose to focus the rest of this study on ACC because of its sensitivity to conspecific distress and its ability to influence behavioral choices.

### ACC^CAMKII^ but not ACC^VGAT^ neurons are differentially activated in dams and sires during pup retrieval

The brain-wide activity mapping experiments were limited by the low temporal resolution of *c-fos*. Therefore, we performed fiber photometry in ACC of freely moving dams and sires during pup retrieval (Figure 3A-B). We expressed GCaMP6s in excitatory neurons (ACC^CAMKII^) by restricting GCaMP expression with the CAMKII promoter in ACC of CBA/CaJ mice (Figure 3C). We also expressed GCaMP7s in inhibitory neurons (ACC^VGAT^) by restricting GCaMP expression in a cre-dependent manner in VGAT-Cre mice (Figure 3D). We recorded calcium activity from PNDs 0 – 5 during interactions with pups. We found peaks of activity in ACC^CAMKII^ neurons locked to pup retrieval in dams and sires that decreased in magnitude over days, measured as area under the curve (AUC) (Figure 3E-H). Additionally, we observed a significant sex difference in the magnitude of these calcium transients, with dams showing a stronger activation of ACC^CAMKII^ neurons during retrieval events compared to sires (Figure 3I-K). The decrease in magnitude of responses over days was not caused by degradation of the fluorescent signal. We expressed GCaMP6s in ACC^CAMKII^ neurons of a separate cohort of females and recorded their activity during interactions with pups from their first and second litters (Figure S5). Activity during pup retrieval was stronger on early postnatal days and disappeared by PND5 in recordings from the first litter; strong activity subsequently reappeared in recordings from a second litter (Figure S5).

**Figure 3.**
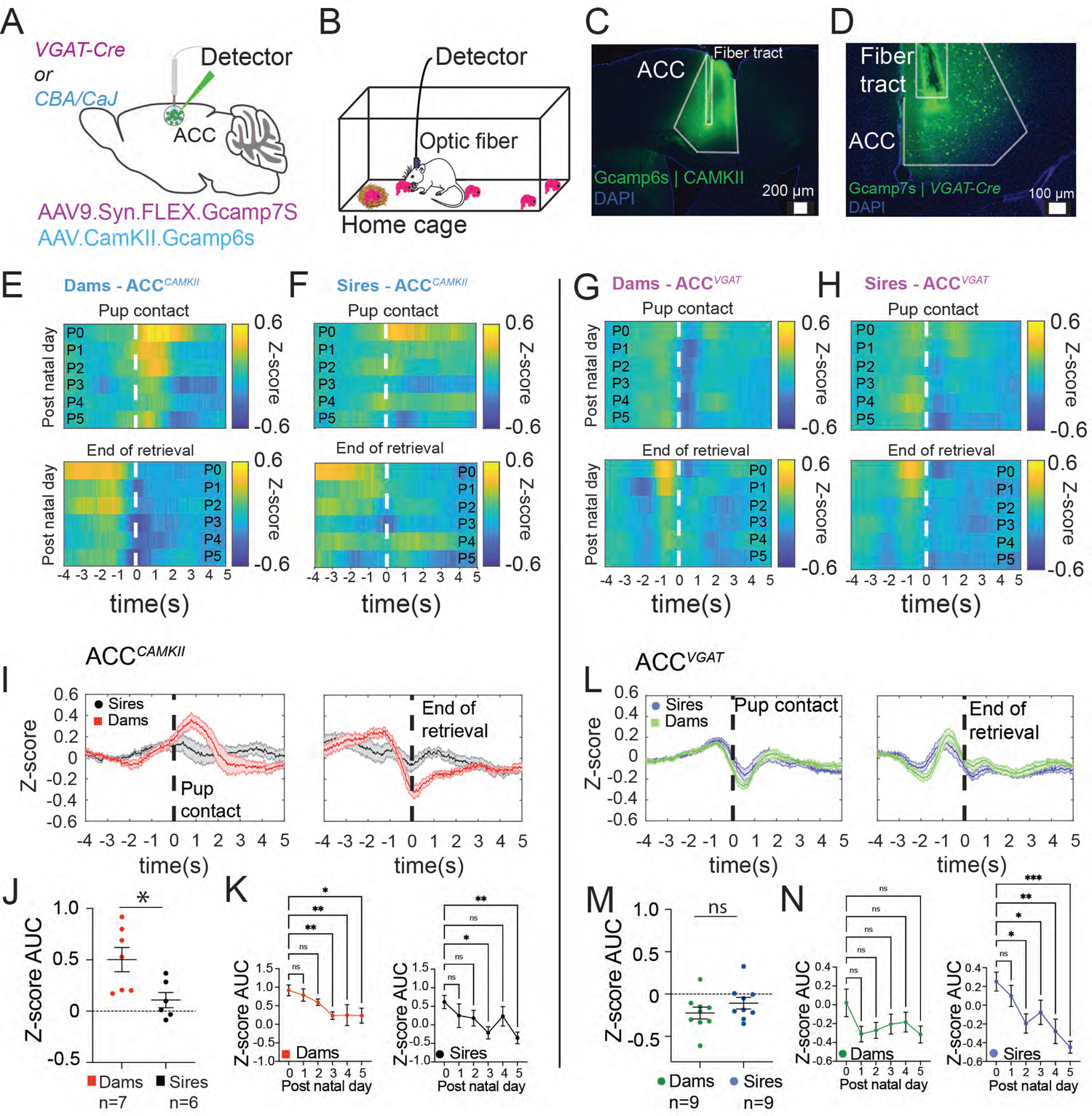
ACC^CAMKII^ but not ACC^VGAT^ neurons are differentially activated in dams and sires during pup retrieval behavior. **A)** Schematic depicting our viral strategy to express GCaMP7s in excitatory and inhibitory neurons in the anterior cingulate cortex of dams and sires. **B)** Schematic of behavioral paradigm. **C)** Representative photomicrograph of a coronal brain section showing fiber placement and GCaMP6s expression in ACC^CAMKII^ neurons. **D)** Representative photomicrograph of a coronal brain section showing fiber placement and GCaMP7s expression in ACC^VGAT^ neurons. **E)** Heatmaps of mean GCaMP6s fiber photometry signals from excitatory neurons in the cingulate cortex during pup gathering events in dams (n = 7). The top panel is a heatmap of data from retrieval events aligned to the pup contact. Each row is the mean activity across all mice for each of 6 days. The bottom panel shows the same signals aligned to the end of the retrieval events. **F)** Heatmaps of mean GCaMP6s fiber photometry signals from excitatory neurons in the cingulate cortex during pup gathering events in sires (n = 6). Panels are arranged as in (E). **G)** Heatmaps of mean GCaMP7s fiber photometry signals from inhibitory neurons in the cingulate cortex during pup gathering events in dams (n = 9). The top panel is a heatmap of data from retrieval events aligned to the pup contact. Each row is the mean activity across all mice for each of 6 days. The bottom panel shows the same signals aligned to the end of the retrieval events. **H)** Heatmaps of mean GCaMP7s fiber photometry signals from inhibitory neurons in the cingulate cortex during pup gathering events in sires (n = 6). Panels are arranged as in (G). **I)** Plots of the mean Z-scored traces of excitatory neurons for all mice and all days, contrasting dams (red) and sires (black). The top panel shows the activity aligned to the pup contact and the bottom panel shows the activity aligned to the end of the retrieval. **J)** Comparison of the mean magnitude of the retrieval-related activity of excitatory neurons between dams and sires, quantified as the mean area under the curve of traces from all days between pup contact and 2 seconds after. (Mann Whitney U test, \**p* = 0.035) **K)** Comparison of the mean magnitude of the retrieval-related activity of excitatory neurons in dams and sires, quantified as the mean area under the curve of traces showing a decline in the magnitude of activity over post-natal days 0 – 5. The left panel shows dam’s responses (Kruskal-Wallis test **p=0.0093; Benjamini, Krieger and Yekutieli multiple comparison test, P0 vs. P3 **p=0.0049; P0 vs. P4 **p=0.0033; P0 vs. P5 *p=0.0209). The right panel shows sire’s responses (Kruskal-Wallis test *p=0.0433; Benjamini, Krieger and Yekutieli multiple comparison test, P0 vs. P3 *p=0.0083; P0 vs. P5 **p=0.0030). **L)** Plots of the mean Z-scored traces of inhibitory neurons for all mice and all days contrasting dams (green) and sires (blue). The top panel shows the activity aligned to pup contact, and the bottom panel shows the activity aligned to the end of the retrieval. **M)** Comparison of the mean magnitude of the retrieval-related activity of inhibitory neurons between dams and sires, quantified as the mean area under the curve of traces from all days between pup contact and 2 seconds after. (Mann Whitney test, *p* =0.3401). **N)** Comparison of the mean magnitude of the retrieval-related activity of inhibitory neurons between dams and sires, quantified as the mean area under the curve of traces showing a decline in the magnitude of activity over post-natal days 0 – 5. The left panel shows dam’s responses (Kruskal-Wallis test not significant p=0.2914). The right panel shows sire’s responses (Kruskal-Wallis test ***p=0.0008; Benjamini, Krieger and Yekutieli multiple comparison test P0 vs. P2 *p= 0.0228; P0 vs. P3 *p=0.0571; P0 vs. P4 **p=0.0020; P0 vs. P5 ***p<0.0001).

In contrast to ACC^CAMKII^ neurons, ACC^VGAT^ neurons showed an abrupt reduction in activity at pup contact followed by a peak of activity just before the end of each retrieval event (Figure 3G-H). In contrast to the ACC^CAMKII^ neurons, we did not observe any significant sex-dependent differences in ACC^VGAT^ neuron activity during pup retrieval behavior (Figure L-N). We confirmed the position of the optical fibers in ACC using immunohistochemistry (Figure S6 and S7).

We also observed sex differences in the relationship between ACC^CAMKII^ and ACC^VGAT^ neurons. The inverse correlation between the two cell types appeared to be weaker in sires compared to dams (Figure 4). In dams, we observed that the activity of ACC^VGAT^ neurons was inversely related to that of ACC^CAMKII^ neurons during pup retrieval behavior (Figure 4 B-C). We measured the magnitude of the responses by quantifying the AUC, and we found that ACC^CAMKII^ neurons increased their activity while ACC^VGAT^ neurons decreased their activity after pup contact (Figure 4D). Additionally, we measured the magnitude of the responses of inhibitory and excitatory populations at the end of the retrieval events and found that ACC^VGAT^ neurons showed stronger activation compared to ACC^CAMKII^ neurons when the dams dropped the pups in the nest (Figure 4E).

**Figure 4.**
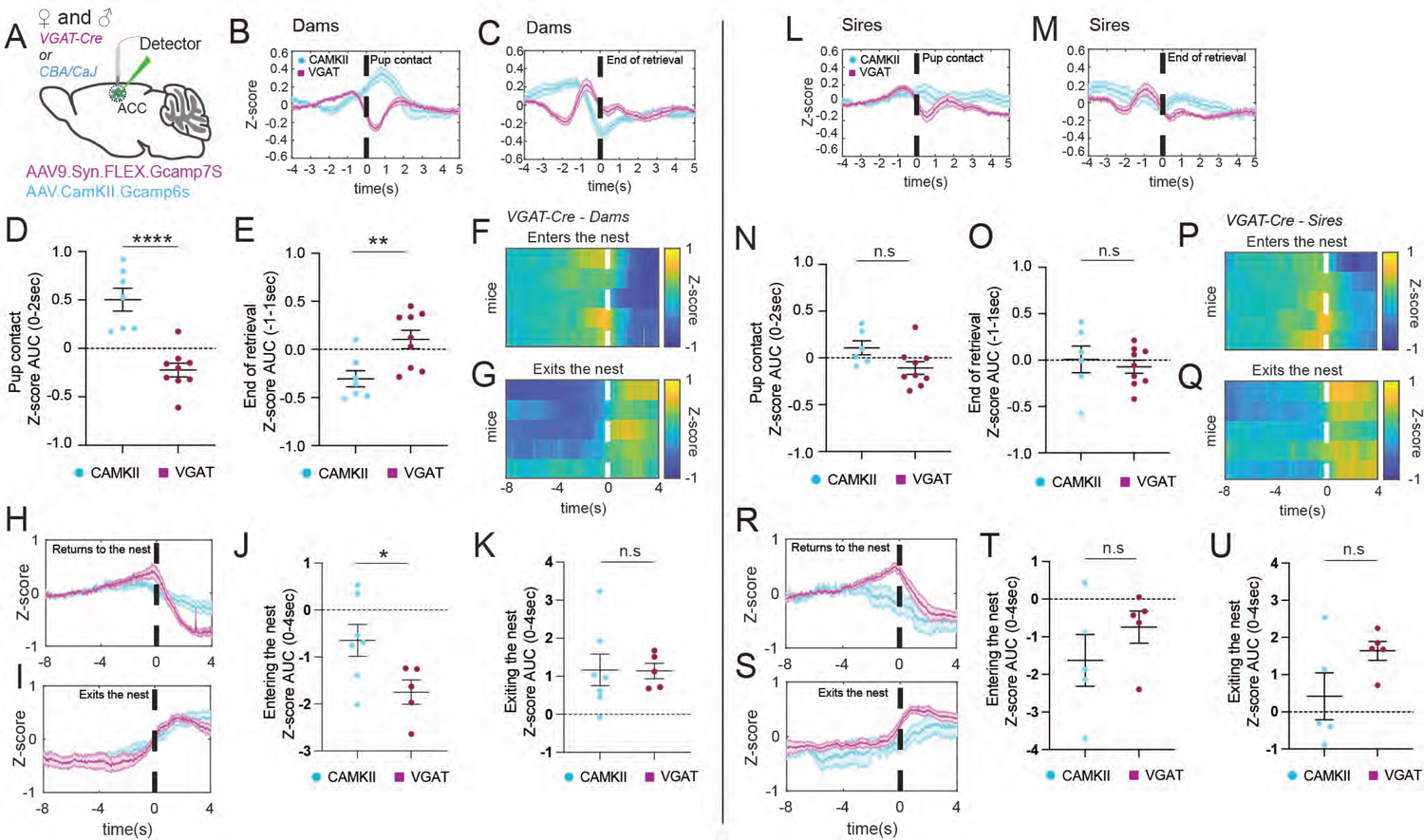
ACC^CAMKII^ and ACC^VGAT^ cell populations show reciprocal activation during pup retrieval behavior in dams but not in sires. **A)** Schematic depicting our viral strategy to express GCaMP7s in ACC^VGAT^ and ACC^CAMKII^ neurons in dams and sires. **B)** Plot of the mean Z-scored traces of excitatory neurons (cyan) and inhibitory neurons (magenta) aligned to pup contact during pup retrieval for all dams and all days (n = 7 mice and n = 9 mice respectively). **C)** Plot of the mean Z-scored traces of excitatory neurons (cyan) and inhibitory neurons (magenta) during pup retrieval for all dams and all days (n = 7 mice and n = 9 mice respectively. **D)** Comparison of the mean magnitude of retrieval-related activity between excitatory neurons and inhibitory neurons in dams, quantified as the mean area under the curve of traces from all days between pup contact and 2 seconds after. (Unpaired t-test ****p<0.0001) **E)** Comparison of the mean magnitude of end of retrieval-related activity between excitatory neurons and inhibitory neurons in dams, quantified as the mean area under the curve of traces from all days between 1 second before and 1 second after the end of retrieval. (Unpaired t-test **p=0.0074). **F-G)** Heatmaps of mean GCaMP7s fiber photometry signals from inhibitory neurons in the cingulate cortex of dams during free interactions with pups. The signals are aligned to the entry of the dam to the nest (F) and the exit of the dam from the nest (G) after retrieval of all pups. Each row represents the mean z-score from postnatal day 0-5 for each mouse (n = 5 mice). **H-I)** v). **J)** Comparison of the mean magnitude of nest entry-related activity between excitatory neurons (cyan) and inhibitory neurons (magenta) in dams, quantified as the mean area under the curve of traces from all days between the dam entering the nest and 4 seconds after. (Unpaired t-test *p=0.0378). **K** Comparison of the mean magnitude of nest exit-related activity between excitatory neurons and inhibitory neurons in dams, quantified as the mean area under the curve of traces from all days between the dam exiting the nest and 4 seconds after. (Unpaired t-test n.s.; p=0.9613). **L)** Plot of the mean Z-scored traces of excitatory neurons (cyan) and inhibitory neurons (magenta) aligned to pup contact during pup retrieval for all sires and all days (n = 6 mice and n = 5 mice respectively). **M)** Plot of the mean Z-scored traces of excitatory neurons (cyan) and inhibitory neurons (magenta) aligned to the end of pup retrieval for all sires and all days (n = 7 mice and n = 9 mice respectively.). **N)** Comparison of the mean magnitude of retrieval-related activity between excitatory neurons and inhibitory neurons in sires, quantified as the mean area under the curve of traces from all days between pup contact and 2 seconds after. (Unpaired t-test, n.s; p=0.0581). **O)** Comparison of the mean magnitude of retrieval-related activity between excitatory neurons and inhibitory neurons in sires, quantified as the mean area under the curve of traces from all days between 1 second before and 1 second after the end of retrieval. (Unpaired t-test, n.s; p=0.5936). **P-Q)** Heatmaps of mean GCaMP7s fiber photometry signals from inhibitory neurons in the cingulate cortex of sires during free interactions with pups. The signals are aligned to the entry of the sire to the nest (P) and the exit of the sire from the nest (Q) after retrieval of all pups. Each row represents the mean z-score from postnatal day 0-5 for each mouse (n = 5 mice). **R-S)** Plots of the mean Z-scored traces of excitatory neurons (cyan) and inhibitory neurons (magenta) for all sires and all days aligned to the sire’s entry to the nest (R) and exit from the nest (S) (n = 5 mice). **T)** Comparison of the mean magnitude of nest entry-related activity between excitatory neurons (cyan) and inhibitory neurons (magenta) in sires, quantified as the mean area under the curve of traces from all days between the dam entering the nest and 4 seconds after. (Unpaired t-test, n,s; p=0.3085). **U)** Comparison of the mean magnitude of nest exit-related activity between excitatory neurons and inhibitory neurons in sires, quantified as the mean area under the curve of traces from all days between the sire exiting the nest and 4 seconds after. Unpaired t-test not significant, n.s; p=0.1092).

ACC^VGAT^ neurons exhibited a sex-dependent decrease in activity when the dam entered the nest after all the pups had been retrieved. Activity increased again to the baseline level when the dam exited the nest (Figure 4F-I). The activity of ACC^VGAT^ and ACC^CAMKII^ neurons was significantly different when the dams entered, but not when they exited the nest (Figure 4J-K). In sires, we did not observe any significant differences between ACC^CAMKII^ and ACC^VGAT^ neurons during parenting (Figure 4L-U). These data argue that there is sex-dependent involvement of ACC in pup retrieval and other parenting behaviors.

### Silencing ACC^CAMKII^ neurons increases parental neglect

Based on these activity patterns during retrieval, we speculated that ACC activity, specifically ACC^CAMKII^ neuron activity, is necessary for attentive parenting in dams and sires. Therefore, we expressed either the inhibitory DREADD (Designer Receptor Exclusively Activated by Designer Drugs) hM4D(Gi) or GFP in ACC^CAMKII^ neurons (Figure 5A) enabling us to silence ACC^CAMKII^ neurons with i.p. injection of clozapine ^51, 52^. We confirmed selective expression of the inhibitory DREADDS in ACC using immunohistochemistry (Figure 5B-C, S8, and S9). All mice were injected with clozapine or saline on alternating days (clozapine on P0/P2 and saline on P1/P3) and ran through the protocol depicted in Figure 5D. Chemogenetic inactivation of ACC^CAMKII^ neurons in dams disrupted pup retrieval behavior in early PNDs compared to GFP controls (Figure 5E-F). These results are consistent with our photometry data showing that ACC^CAMKII^ neurons are more strongly activated during pup retrieval in early PNDs (0-3) compared to later PNDs (4-5). These observations suggest that ACC modulates retrieval behavior, particularly in the first few days after birth.

**Figure 5.**
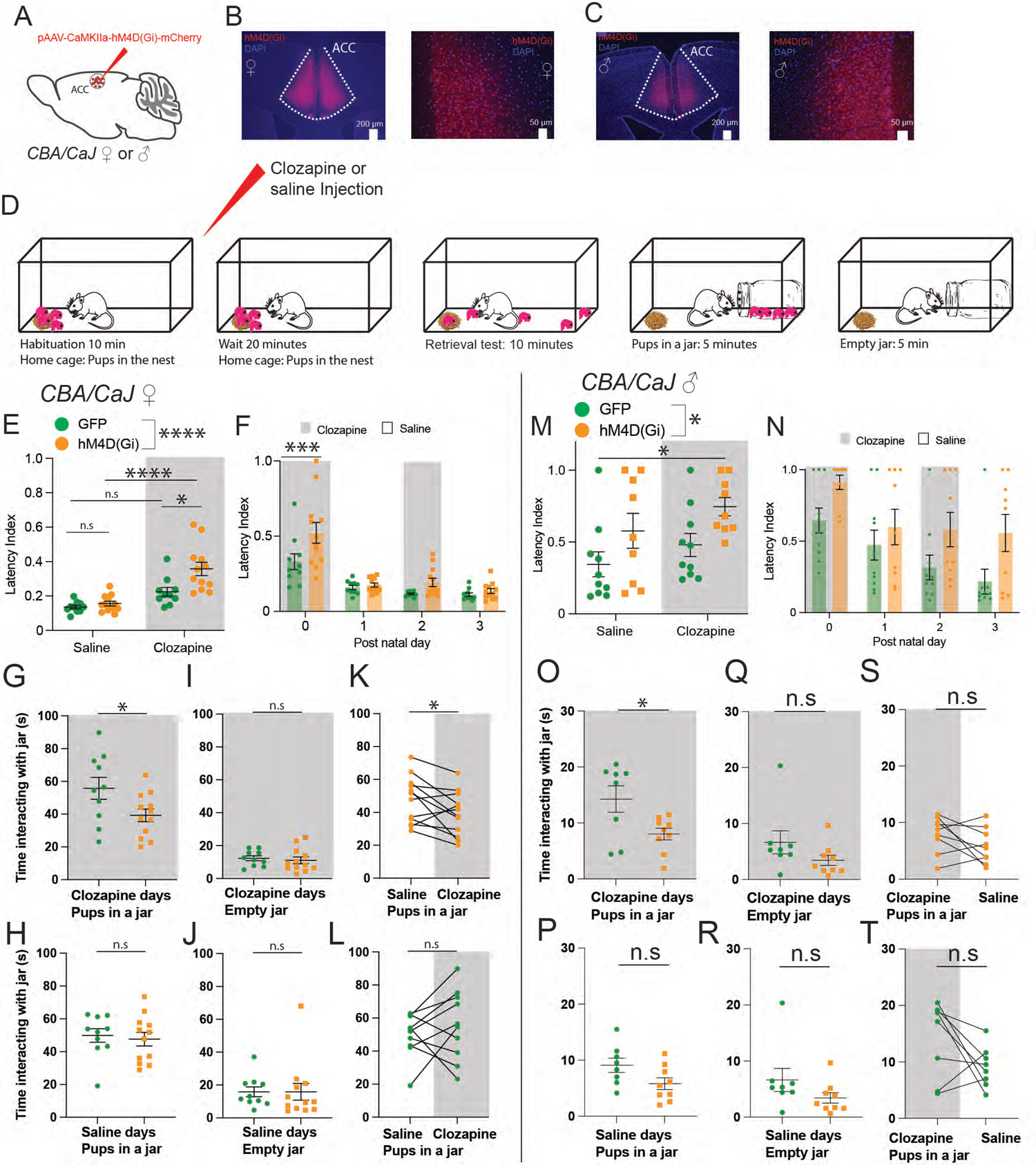
Chemogenetic inactivation of ACC^CAMKII^ neurons disrupts pup-directed behaviors. **A)** Schematic of the viral strategy we used to inactivate ACC^CAMKII^ neurons in dams and sires during interactions with pups. **B-C)** Representative photomicrographs of the expression of the hM4D(Gi) construct in the ACC of dams and sires. **D)** Schematic of the behavioral paradigm used to measure pup retrieval performance and time interacting with pups in a jar. **E)** Scatterplot of mean latency index (± s.e.m.) in GFP expressing dams injected with saline (n = 10), GFP expressing dams (green) injected with clozapine (n = same 10 mice from saline experiment), hM4D(Gi) expressing dams (orange) injected with saline (n = 12), and hM4D(Gi) expressing dams injected with clozapine (n = same 12 mice from saline experiment); Ordinary one-way ANOVA; **p<0.0001; F=15.98; Tukey’s multiple comparison test, Clozapine: GFP vs. Clozapine: hM4D(Gi) **p=0.0039, Saline: hM4D(Gi) vs. clozapine: hM4D(Gi) ****p<0.0001. **F)** Scatterplot of the same data as in E, separated by different post-natal days. Two-way ANOVA with main effects for day (****p<0.0001) and GFP vs DREADDS (**p=0.0025) and no significant interaction p=0.05; Sidak’s multiple comparison test, ***p=0.0009 for P0; P1-P3: n.s. p>0.05). **G)** Plot comparing time spent interacting with the jar of pups for GFP-expressing dams (green; n = 10) and hM4D(Gi) expressing dams (orange; n = 12) injected with clozapine (Mann Whitney test; \**p* = 0.0426). **H)** Plot comparing time spent interacting with the jar of pups for GFP-expressing dams (green; n = 10) and hM4D(Gi) expressing dams (orange; n = 12) injected with saline (Mann Whitney test p=0.6277). **I)** Plot comparing time spent interacting with the empty jar for GFP-expressing dams (green; n = 10) and hM4D(Gi) expressing dams (orange; n = 12) injected with clozapine (Mann Whitney test p=0.4176). **J)** Plot comparing time spent interacting with the empty jar for GFP-expressing dams (green; n = 10) and hM4D(Gi) expressing dams (orange; n = 12) injected with saline (Mann Whitney test p=0.3463). **K)** Plot comparing time spent interacting with the jar of pups by hM4D(Gi) expressing dams injected with saline (n = 12) and hM4D(Gi) expressing dams (n = 12) injected with clozapine (Paired t test; *p=0.0369). **L)** Plot comparing time spent interacting with the empty jar by GFP expressing dams injected with saline (n = 10) and GFP expressing dams (n = 10) injected with clozapine (Paired t test; p=0.4038). **M)** Scatterplot of mean latency index (± s.e.m.) in GFP expressing sires injected with saline (n = 10), GFP expressing sires (green) injected with clozapine (n = same 10 mice from saline experiment), hM4D(Gi) expressing sires (orange) injected with saline (n = 9), and hM4D(Gi) expressing sires injected with clozapine (n = same 9 mice from saline experiment); Ordinary one-way ANOVA *p=0.02; F=3.521; Tukey’s multiple comparison test, Saline: GFP vs. clozapine: hM4D(Gi) *p=0.0168, no other significant comparisons. **N)** Scatterplot of the same data as in M, but separated by different post-natal days. Two-way ANOVA with main effects for day (**p=0.0014) and GFP vs DREADDS (***p=0.0008) and no significant interaction p=0.7601; Sidak’s multiple comparison test not significant. **O)** Plot comparing time spent interacting with the jar of pups for GFP-expressing sires (n = 8) and hM4D(Gi) expressing sires (n = 9) injected with clozapine (Mann Whitney test p=0.0592). **P)** Plot comparing time spent interacting with the jar of pups for GFP-expressing sires (n = 8) and hM4D(Gi) expressing sires (n = 9) injected with saline (Mann Whitney test not significant). **Q)** Plot comparing time spent interacting with the empty jar for GFP-expressing sires (n = 8) and hM4D(Gi) expressing sires (n = 9) injected with clozapine (Mann Whitney test not significant). **R)** Plot comparing time spent interacting with the empty jar for GFP-expressing sires (n = 8) and hM4D(Gi) expressing sires (n = 9) injected with saline (Mann Whitney test p=0.2660). **S)** Plot comparing time spent interacting with the empty jar for GFP-expressing sires injected with saline (n = 9) and hM4D(Gi) expressing sires (n = 9) injected with clozapine (Paired t test p=0.1392). **T)** Plot comparing time spent interacting with the empty jar for GFP-expressing sires injected with saline (n = 8) and GFP expressing sires (n = 8) injected with clozapine (Paired t test p=0.1024).

In the same group of mice, we recorded interactions with the pups trapped in a jar or an empty jar 30 minutes after clozapine or saline injections (Figure 5D, Supplementary Movie 1). hM4D(Gi)-expressing dams spent significantly less time interacting with the jar relative to GFP controls after clozapine injection (Figure 5G). On days when clozapine was injected, hM4D(Gi)-expressing dams spent significantly less time interacting with the jar as compared to days when saline was injected (Figure 5K). There was no difference in this behavior between the hM4D(Gi) - and GFP-expressing dams when they were injected with saline (Figure 5H). No differences were observed between any of the groups in their behavior in the presence of an empty jar (Figure 5I-J, L). These results show that silencing ACC^CAMKII^ neurons in dams increases pup neglect and may also decrease maternal motivation.

We used the same strategy to assess the consequences of inactivating ACC^CAMKII^ neurons in sires. We found that chemogenetic inactivation of ACC^CAMKII^ neurons in sires does not disrupt pup retrieval significantly (Figure 5M-N). However, inactivation of ACC^CAMKII^ neurons on PND0 affected the sires’ pup retrieval performance on subsequent days such that hM4D(Gi)-expressing animals appear to be impaired even after a saline injection (Figure 5M-N). With regard to pups trapped in the jar, we found that all sires spent significantly less time than dams interacting with in a jar containing pups, but they did not differ from dams in their interactions with an empty jar. hM4D(Gi)-expressing sires spent significantly less time interacting with the pups in the jar than GFP controls when injected with clozapine (Figure 5O), but this difference was not observed when injected with saline (Figure 5P). We did not see any significant differences in the time spent interacting with an empty jar between hM4D(Gi) expressing mice and GFP controls injected with clozapine or saline (Figure 5Q-R). Chemogenetic inactivation of ACC^CAMKII^ neurons did not affect sires’ behavior when comparing hM4D(Gi)-expressing mice injected with clozapine or saline (Figure 5S). Collectively, our chemogenetic data show that inactivation of ACC^CAMKII^ neurons increases pup neglect and decreases sensitivity to respond to pup distress in dams and not in sires.

Importantly, the effect on behavior that we observed from inactivating ACC^CAMKII^ neurons was not due to a direct effect on anxiety-like behaviors. We tested hM4D(Gi)-expressing and GFP-expressing dams and sires injected with clozapine on an elevated-plus maze (Figure S10A-B). We found no significant differences in the percent of time animals spent in the open arms (Figure S10C and E) or the number of entries to the open arms (Figure S10D and F).

### Sex-dependent signaling from LC to ACC during pup retrieval

Previous reports have described the afferents ^29^ and efferents ^53^ of ACC. Interestingly, ACC is interconnected with the noradrenergic locus coeruleus (LC) ^29, 53, 54^, and activity patterns in ACC and LC are coordinated ^30^. Previously we proposed, based on recordings from female mice, that LC contributes to goal-directed action selection during parental behavior with global release of noradrenaline (NA) ^26^. However, the downstream targets of LC that modulate social behavior according to NA signaling remain unknown. Therefore, we injected a retrograde adeno-associated virus (AAV) driving expression of the fluorescent reporter tdTomato in ACC and confirmed the existence of a projection from LC to ACC (Figure 6A).

**Figure 6.**
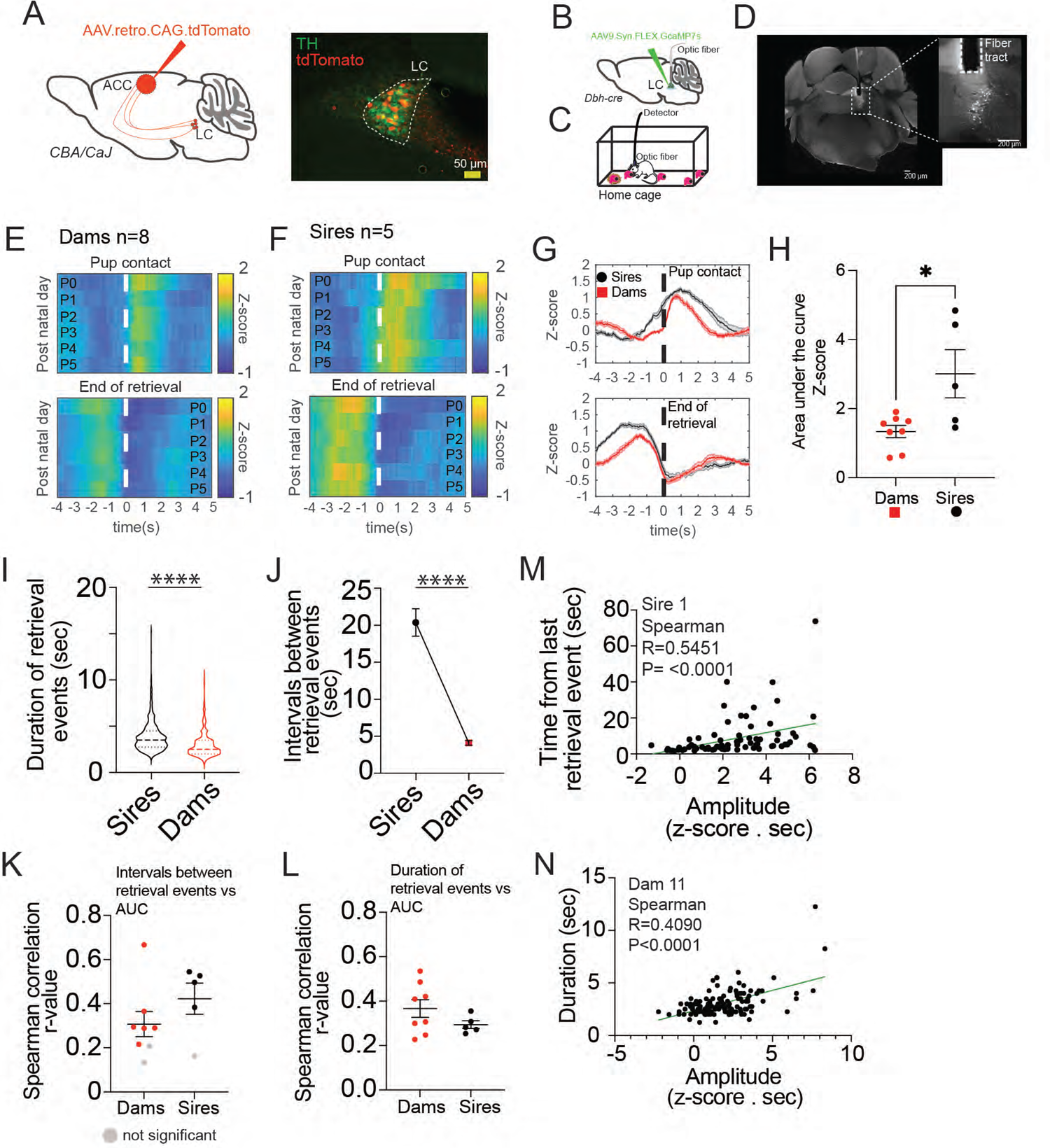
Timing and magnitude of LC activity associated with pup retrieval behavior is different in sires compared to dams. **A)** Schematic of virus injection (*left*) and a representative photomicrograph of a coronal brain section showing retrograde labeling from ACC to LC (*right*). Green shows tyrosine hydroxylase (TH) antibody staining and red shows tdTomato expression driven by the retrograde AAV injection. **B)** Schematic of the viral strategy we used to selectively express GCaMP7s in LC^DBH^ neurons. **C)** Schematic of the pup retrieval behavior paradigm. **D)** Representative photomicrograph of a coronal brain section showing GCaMP7s expression in the LC. The placement of the optical fiber is also visible (*inset*). **E-F)** Heatmaps of mean GCaMP7s fiber photometry signals from LC during pup gathering events in dams (E) (n = 8 mice) and sires (F) (n = 5) The top panel is a heatmap of data from retrieval events aligned to the pup contact. Each row is the mean activity across all mice for each of 6 days. The bottom panel shows the same signals aligned to the end of the retrieval events. **G)** Plots of the mean Z-scored traces of LC fiber photometry signals for all mice and all days, contrasting dams (red) and sires (black). The top panel shows the activity aligned to the pup contact and the bottom panel shows the activity aligned to the end of the retrieval. **H)** Comparison of the mean magnitude of the retrieval-related activity of inhibitory neurons between dams and sires, quantified as the mean area under the curve of traces from all days between pup contact and 4 seconds after. (Mann-Whitney test, *p* = 0.0451). **I)** Violin plots comparing the duration of retrieval events between dams and sires (sires n = 5 mice, 618 events; dams n = 8 mice, 1238 events; Mann-Whitney test p < 0.0001). **J)** Duration of intervals in between retrieval events (sires n = 5 mice, n = 508 events; dams n = 8 mice, n = 1072 event; Mann-Whitney test p < 0.0001). **K)** Scatter plot of coefficients (r) of Spearman correlation of the intervals between retrieval events and the magnitude of LC responses during pup retrieval in dams and sires. Gray dots represent correlations that did not reach significance. **L)** Scatter plot of coefficients (r) of Spearman correlation of the duration of the retrieval events and the magnitude of LC responses during pup retrieval in dams and sires. Gray dots represent correlations that did not reach significance. **M)** Example scatterplot of the time between retrieval events and the magnitude of LC responses in a sire. The green line represents a linear regression. **N)** Example scatterplot of the duration of retrieval events and the magnitude of LC responses in a dam. The green line represents a linear regression.

We compared the LC activity associated with pup retrieval in sires to that in dams by performing fiber photometry of GCaMP7s signal in LC of freely behaving mice. We injected a Cre-dependent AAV virus driving expression of the fluorescent Ca^2+^ reporter GCaMP7s into DBH-Cre mice expressing Cre recombinase in cells that produce dopamine beta-hydroxylase. This enzyme performs the final step in the synthesis of noradrenaline, and therefore our injections selectively labeled noradrenergic LC neurons (Figure 6B-C). LC responses associated with pup retrieval were longer in sires compared to dams (Figure 6E-G). These responses were sustained for the entire pup retrieval event and returned to baseline activity levels when the mouse drops the pup in the nest (Figure 6G). By measure of the AUC extending 4 s beyond pup contact, LC responses were weaker in dams compared to sires (Figure 6H). Over all days, retrieval events were longer in sires (3.77 ± 0.04 s) than in dams (3.15 ± 0.03 s) (Figure 6I), and sires had significantly longer intervals between retrieval events (sires: 20.37 ± 1.86 s; dams: 4.1 ± 0.31 s) (Figure 6J). Not surprisingly, the magnitude of LC responses was positively correlated with the duration of retrieval events (Figure 6L-N). Interestingly, the magnitude of the LC response and the time since the preceding retrieval event were also positively correlated (Figure 6K-M). We conclude that the temporal precision of the neural activity in LC reflects sex differences in the temporal precision of pup retrieval.

The high amplitude of phasic activity in LC during pup retrieval implies that the activity is pervasive through most LC neurons ^26^. We found that neurons in LC that project to ACC (LC-ACC) share this phasic activity pattern. We injected a retrograde AAV in ACC to express Cre recombinase in LC-ACC, and we injected Cre-dependent GCaMP7s AAV in LC (Figure 7A-B). Indeed, during pup retrieval (Figure 7C), we found that LC-ACC neurons exhibited significant, temporally precise calcium transients time-locked to pup contact that were similar to those seen from all neurons in LC (Figure 7D,E). These events very likely resulted in NA release in the ACC during pup retrieval. We injected an NA sensor ^55^ in ACC, and we observed NA release associated with pup retrieval (Figure 7G-I). Comparing the AUC of the NA signal at baseline to the AUC after pup contact, there was a significant rise in NA release during pup retrieval relative to baseline (Figure 7J). These data establish a functional connection between LC and ACC associated with parental behavior.

**Figure 7.**
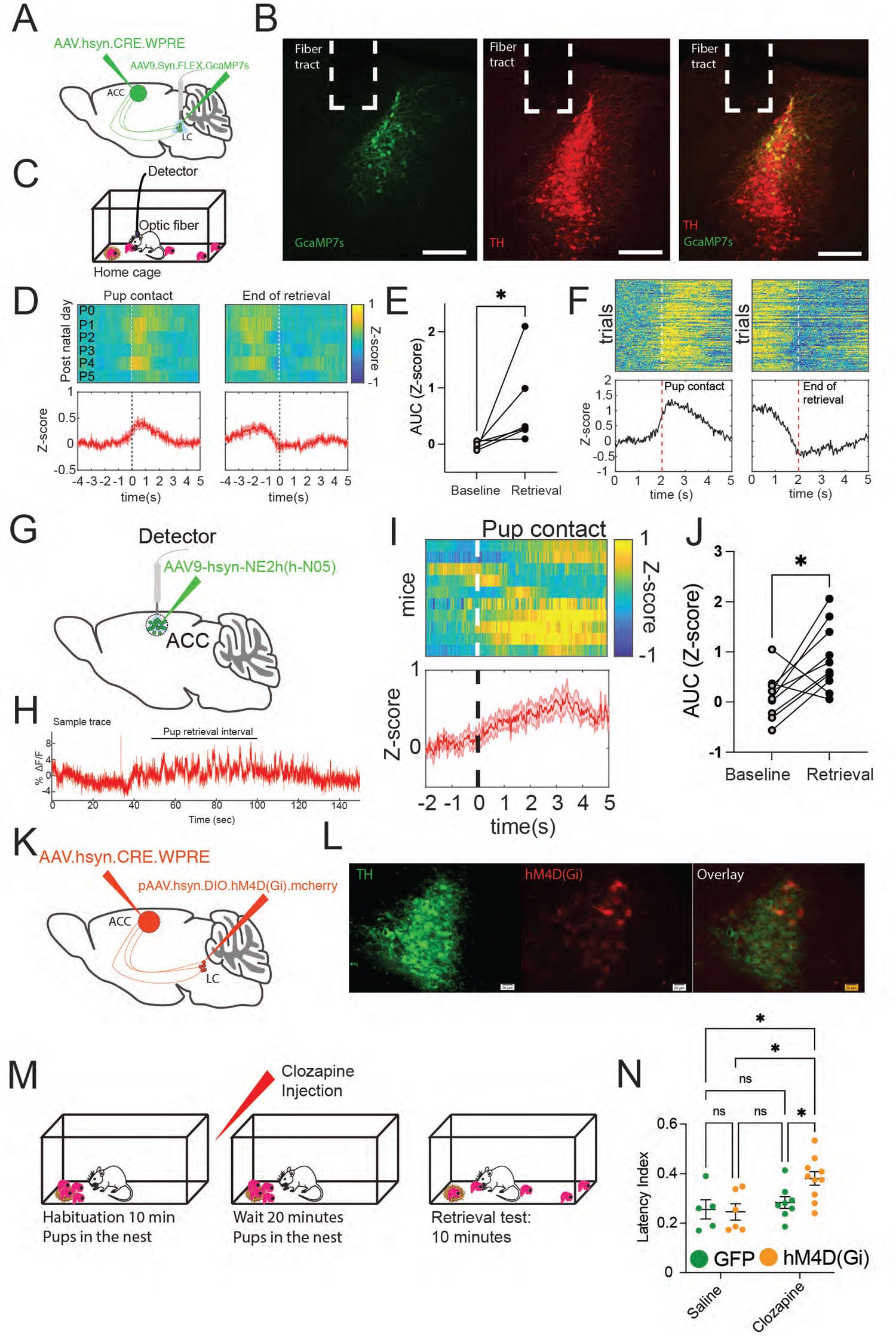
LC-ACC neurons are active during pup retrieval behavior, release noradrenaline in ACC, and selective inactivation of LC-ACC neurons disrupts pup retrieval behavior. **A)** Schematic of the viral strategy we used to express GCaMP7s in LC neurons that project to the ACC. **B)** Representative photomicrographs of a coronal brain section showing GCaMP7s expression and fiber placement in LC. Green labeling is GCaMP7s and red labeling is TH antibody staining. **C)** Schematic of pup retrieval behavior. **D)** Heatmaps and plots of mean Z-scored traces for LC-ACC neurons during pup retrieval. The left panel shows a heatmap (top) in which each row is the mean activity across all mice for each of 6 days and a plot (bottom) of mean Z-scored GCaMP7s traces from LC-ACC neurons for all mice and all days. Data are aligned to the pup contact for each retrieval event. The right panel is the same data but aligned to the end of retrieval events. **E)** Plot of the mean magnitude of the retrieval-related activity of LC-ACC neurons quantified as the mean area under the curve between pup contact and 2 seconds after, compared with baseline activity (Wilcoxon matched-pairs signed rank test *p=0.0312). **F)** Heatmaps and plots of mean GCaMP7s fiber photometry signals from LC-ACC neurons during pup gathering events in one representative dam. The left panel shows a heatmap (top) in which each row is the activity from one retrieval trial, and a plot (bottom) of mean Z-scored GCaMP7s traces from LC-ACC neurons for all days. Data are aligned to the pup contact for each retrieval event. The right panel is the same data but aligned to the end of retrieval events. **G)** Schematic of our viral strategy to express a noradrenaline sensor in ACC. **H)** Representative dF/F trace of fluorescence detected noradrenaline release as a dam interacted with pups. The black line above the trace indicates the time period when the dam was retrieving the pups to the nest. **I)** Heatmap and plot of mean Z-scored fluorescent GRAB_NE_ signal from ACC during pup gathering events in dams (n=6) and sires (n=4). The top panel is a heatmap in which each row is the mean activity across all sessions from one mouse. The bottom panel shows the mean Z-scored fluorescence trace for all mice. Data are aligned to the pup contact for each retrieval event. **J)** Plot of the mean magnitude of the retrieval-related activity of GRAB_NE_ signal from ACC during pup gathering events quantified as the area under the curve for the NA release responses from the pup contact to 2 seconds after and 2 seconds of baseline (Paired t test *p=0.0173). **K)** Schematic of the viral strategy we used to express inhibitory DREADDS only in LC neurons that project to the ACC. **L)** Representative photomicrograph of a coronal brain section showing DREADDS expression in LC neurons that project to ACC. Green labeling is tyrosine hydroxylase (TH) antibody staining and red is hM4D(Gi) expression. **M)** Schematic of the behavioral paradigm used to measure pup retrieval performance. **N)** Scatterplot of mean latency index (± s.e.m.) in GFP expressing dams injected with saline (n = 10), GFP expressing dams (green) injected with clozapine (n = same 10 mice from saline experiment), hM4D(Gi) expressing dams (orange) injected with saline (n = 9), and hM4D(Gi) expressing dams injected with clozapine (n = same 9 mice from saline experiment); Ordinary one-way ANOVA; **p=0.0075; F=5.042; Benjamini, Krieger and Yekutieli multiple comparison test, Clozapine: GFP vs. Clozapine: hM4D(Gi) *p=0.0137, Saline: hM4D(Gi) vs. clozapine: hM4D(Gi) **p<0.0028, Saline: GFP vs. Clozapine: hM4D(Gi) *p=0.0072.

### Selectively silencing noradrenergic input to the ACC impairs pup retrieval

Finally, we tested whether the LC-ACC circuit is necessary for parental behavior. We injected a retrograde AAV virus in ACC to express Cre recombinase in LC-ACC neurons and Cre-dependent hM4D(Gi) in LC (Figure 7K-L). We recorded interactions with pups 20 minutes after clozapine injection starting on PND 0 – 3 (Figure 7M). We found that chemogenetic inactivation of LC-ACC neurons disrupts pup retrieval behavior in early PNDs (Figure 7N). Although the inactivation of the LC-ACC circuit did not abolish pup retrieval behavior completely, it increased pup neglect and the latency to retrieve pups in dams.

## DISCUSSION

Empathy can be defined as the adoption of another individual’s emotional state. In the past few decades, rodents have been shown to display empathy-like behaviors such as vicarious fear learning ^7^, social transfer of pain and analgesia ^10^, and pro-social behaviors such as consolation of others ^8, 11^. ACC has been proposed as a hub for information about the emotional state of others ^7, 10^. Here, we propose that the involvement of ACC in parental behavior presents a tractable model to reveal the neural computations that underlie decisions influenced by the social perception of distress. During parental encounters, adults need to process offspring cues, including distress cues, and then select an appropriate behavioral response. We propose that ACC, in coordination with the noradrenergic LC, integrates distress signals from the offspring to promote a behavioral response to a distressed individual.

We assessed whether dams and sires display differential sensitivity to pup distress. Not surprisingly, our data show that paternal behavior is more variable and less robust than maternal behavior, consistent with previous reports. For instance, paternal but not maternal behavior is disrupted by the loss of oxytocin ^56^, and disrupting prolactin signaling in the medial preoptic nucleus impairs paternal behavior ^34^. One possibility for the observed behavioral variability is that males are more sensitive than females to contextual changes ^31^. We observed that paternal behavior is very sensitive to changes in context, and we showed that most males did not retrieve pups in a novel environment. Performing the behavior in a novel context did not significantly alter the dam’s behavior. These results suggest that paternal behavior is subject to additional contextual regulation as compared to maternal behavior and suggest differential sensitivity to pup distress in dams and sires.

The neural substrates that modulate behavioral choices during pro-social behaviors remain elusive. Starting from an unbiased brain-wide activity screen, we associated ACC with parental behavior (Figure 2). We also observed sex-dependent ACC activation patterns in dams and sires that retrieve pups to the nest. Consistent with the behavioral variability observed in sires, *c-fos* expression levels were more variable in sires compared to dams. However, two limitations of immediate early gene screens are that they only provide a snapshot of brain activity from a given time point and that they lack cell type specificity. Therefore, we chose to observe neural activity in freely moving mice to measure the temporal dynamics of ACC neurons during parenting behaviors. We were also interested in whether different cell types in ACC modulate different aspects of pup retrieval behavior. To address those limitations, we assessed the activity dynamics of inhibitory and excitatory neurons in ACC during free interactions with pups.

Consistent with the higher expression levels of *c-fos* in ACC in dams compared to sires, our photometry results showed stronger activation of ACC^CAMKII^ neurons in dams as compared to sires during pup retrieval (Figure 3). Neural responses in ACC during pup retrieval were stronger in early PNDs, presumably when the pups are more vulnerable to environmental conditions, and the behavior is being actively learned. We observe that the responses during pup retrieval start before the animals contact a pup prior to retrieving it to the nest. These results suggest that the ACC^CAMKII^ neural responses are likely associated with the late pre-execution stages of pup retrieval. We believe that ACC may be involved in processing pup distress to influence engagement with the pups. It is possible that some input(s) to ACC may put the animal in a vigilant state during interactions with pups in distress by slowly affecting the excitability of ACC neurons, raising or lowering the threshold for making a decision. The differences in ACC^CAMKII^ neural activity between dams and sires may reflect differential sensitivity to pup distress.

The interplay between inhibitory and excitatory neural populations is essential for nearly all cortical processing ^57^. We recorded the dynamics of inhibitory and excitatory neurons in ACC during pup retrieval using fiber photometry (Figure 4). Interestingly, we observed an inverse relationship between ACC^CAMKII^ and ACC^VGAT^ cells in dams but not in sires. These data suggest that synaptic interactions between excitatory and inhibitory neurons in ACC may be weaker in sires compared to dams. We also observe that inhibitory neurons in ACC exhibit an abrupt decrease in activity when the mice enter the nest with pups, and rapidly return to baseline when the mice exit. It is unclear what sensory attributes of the nest trigger this response. We revealed sex-dependent involvement of inhibitory and excitatory ACC neurons in different components of parental behavior. The role of ACC in parental decision-making lacks a detailed description of the contribution of different cell types, so an open question from this work is to investigate the contribution of different types of excitatory (e.g. Fezf2, PlexinD1), inhibitory (e.g PV, SOM, VIP), or projection neurons in ACC during free interactions with pups.

Inactivation of ACC impairs the acquisition of fear responses by observation ^7^, it also disrupts processing of pain in conspecifics ^10^, and in rats, it disturbs maternal behavior ^15^. We hypothesized that ACC processes distress signals from the pups, and we predicted that inactivating ACC^CAMKII^ neurons may disrupt retrieval. Indeed, chemogenetic inactivation of ACC^CAMKII^ neurons increased the latency to retrieve pups in early PNDs, but the animals’ performance recovered as the pups got older (Figure 5). This is consistent with our photometry data that shows stronger ACC activation during pup retrieval in early PNDs (Figure 3). We speculate that when the pups are younger, mice are more vigilant as a result of the pups being more susceptible to environmental distress.

ACC receives inputs from many brain regions including the noradrenergic LC ^28, 29, 54^. LC plays an important role in arousal, memory formation and retrieval, stress, attention, and goal-directed action selection among many other functions (reviewed in ^58, 59^). However, the neural mechanisms by which the noradrenergic system regulates socially-motivated behavior and social distress remain poorly understood. LC is known to modulate stress responses through corticotropin releasing factor (CRF) which increases tonic firing in LC ^18^. Interestingly, there is sex-dependent sensitivity to CRF in LC neurons, with female LC neurons being more sensitive to CRF compared to male LC neurons ^60^. Thus, LC is a candidate region to modulate sex-dependent stress responses through its projections to ACC and influence parental behavior in response to pup distress. We observed neural responses in LC in dams and sires during pup retrieval behavior (Figure 6). These responses were longer in sires compared to dams and reflected the temporal precision of pup retrieval behavior. These data suggest that sexually divergent activation of LC contributes to sex differences in parental behavior.

We found that the activity patterns in LC-ACC neurons during retrieval closely resembled those from LC-wide recordings. This is consistent with our model that bursts of activity in LC during pup retrieval are pervasive. Pup retrieval also evokes transient NA release in ACC, establishing a functional noradrenergic connection between LC and ACC during parental behavior. Indeed, when we selectively inactivated the LC-ACC, pup retrieval behavior was disrupted in early PNDs consistent with the results from inactivating all ACC^CAMKII^ (Figure 5). Based on our data as a whole, we conclude that ACC maintains sex-dependent sensitivity to pup distress in coordination with the noradrenergic system, and we propose that parental behavior constitutes a tractable model to reveal the neural mechanisms by which social perception of distress can influence decisions.

## ACKNOWLEDGMENTS

The authors would like to thank R. Shansky, J. Tollkühn, H. Hou, and members of the Shea Lab for helpful comments and discussion. This work was supported by grants to SDS from the National Institute of Mental Health (R01MH119250), the C.M. Robertson Foundation, and the Feil Foundation.

## AUTHOR CONTRIBUTIONS

Conceptualization: A.C. and S.D.S., Methodology: A.C., P.O., R.M.-C., and S.D.S. Software: A.C., Formal Analysis: A.C. and R.M.-C. Investigation: A.C., J.C., and R.M.-C. Writing – Original Draft: A.C. and S.D.S. Writing – Review and Editing: A.C. and S.D.S. Visualization: A.C. and S.D.S. Supervision and Funding Acquisition: S.D.S.

## MATERIALS AND METHODS

### Animals

Adult mice (8–14 weeks old) were maintained on a 12h/12 h light-dark cycle (lights on 07:00 h) and received food ad libitum. Genotypes used were CBA/CaJ, VGAT-Cre (C57BL/6 background), and DBH-Cre (Tg(Dbh-cre)KH212Gsat/Mmucd, unfrozen stock, MMRRC). All mice used for pup retrieval experiments were primiparous. All procedures were conducted in accordance with the National Institutes of Health’s Guide for the care and use of laboratory animals and approved by the Cold Spring Harbor Laboratory Institutional Animal Care and Use Committee.

### Pup retrieval assay

The pup retrieval assay was performed as described in ^35^. Briefly, (1) The test subject was habituated with 5 pups in the nest of the home cage for 5 minutes in a soundproof behavioral box, (2) the pups were removed from the cage for 2 minutes, and (3) then the pups were scattered in the cage. The first pup was placed in the nest and then moving clockwise, a pup was placed in each corner and one in the center. Each test subject had a maximum of 5 minutes to gather the pups to the nest. The same procedure was repeated on postnatal days 0 to 5. All assays were performed in the dark during the light cycle and videos were recorded for further analysis.

For the behavioral analysis, we calculated the latency index for each mouse to gather all pups using the formula:

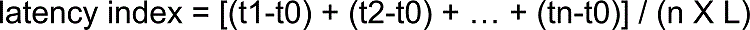

where: n = # of pups outside the nest, t0 = start of trial, tn= time of nth pup gathered, L = trial length.

The same experiment was conducted in a clean/novel cage with different mice.

### Pups in a jar experiment

All pups from the litter were placed in a 4oz. glass jar with a plastic lid with holes in it (AOZITA; B07VHBX3ZC). The test subjects were able to hear and smell the pups in the jar, but were not able to touch them. The animal’s behavior was recorded for 5 min with the pups in the jar and 5 minutes with an empty jar in the home cage. Interactions with the lid of the jar were quantified as a proxy of motivation for the test subject to interact with the pups. All behaviors were scored with the software BORIS ^61^. An interaction with the jar was quantified when the test subject was in close proximity to the lid of the jar and facing it, either touching, biting, or sniffing it.

### Pup USVs recordings

To confirm the jar was enough to put the pups in distress, we recorded USVs for 5 min when the pups were in the nest in the home cage and 5 minutes with the pups in a jar. All pups used to record USVs were 0 – 5 days old. We placed the pups in a jar and placed the jar in the home cage. We started recording the vocalizations using a USV microphone (Condenser ultrasound microphone Avisoft-Bioacoustics CM16/CMPA-5V; part # 40013) 1 minute after placing the pups in the jar. To record the vocalizations in the nest, we removed the parents from the home cage. We started recording USVs 1 minute after removing the parents.

### Behavioral assay for brain-wide *c-fos* induction

Male and female wild-type CBA/CaJ mice breeding pairs were made at 8-10 weeks-old. The experiment included four behavioral groups, and it was performed on post-natal day 3. Baseline: the test subject was kept in the home cage with the pups in the nest for a 3-hour period. Isolated: the test subject was kept in the home cage without the pups for a 3-hour period. Reunion: the test subject was kept in the home cage without the pups for a 90-minute period, and then all pups were returned in the nest for a second 90-minute period. Retrieval: the test subject was kept in the home cage without the pups for 90 minutes, and then all pups were returned scattered in the cage for 90 minutes. The experiment was performed in dams and sires 12-14 weeks old. All experiments were performed in the dark during the light cycle in a sound proof behavioral box. All mice were perfused immediately after the experiment was done through the ascending aorta with 1% PBS, followed by 4% paraformaldehyde (PFA) in 0.1 M PBS (pH 7.4). The brains were removed and post-fixed in paraformaldehyde overnight before starting the clearing protocol.

### Clearing protocol/*c-fos* staining

All brains were cleared using the iDisco+ protocol as described in ^36^.

### Lightsheet imaging

Cleared samples were imaged in sagittal orientation (left hemisphere) on a light-sheet fluorescence microscope (Ultramicroscope II, LaVision Biotec) equipped with a sCMOS camera (Andor Neo) and a 4x/0.5 objective lens (MVPLAPO 4x) equipped with a 6-mm working distance dipping cap. Version v144 of the Imspector Microscope controller software was used. The samples were scanned with a step-size of 3 micrometers using the continuous light-sheet scanning method with the included contrast blending algorithm for the 640 nm and 595 nm channels (20 acquisitions per plane), and without horizontal scanning for the 480-nm channel.

### Statistical analysis for *c-fos* mapping

Statistical comparisons between different groups were run based on either ROIs or evenly spaced voxels. Voxels are overlapping 3D spheres with 100 μm diameter each and spaced 20 μm apart from each other. The cell counts at a given location, Y, are assumed to follow a negative binomial distribution whose mean is linearly related to one or more experimental conditions, X: E[Y]=α+βX. For example, when testing an experimental group versus a control group, the X is a single column showing the categorical classification of mouse sample to group id, i.e. 0 for the control group and 1 for the experimental group ^62, 63^. We found the maximum likelihood coefficients α and β through iterative reweighted least squares, obtaining estimates for sample standard deviations in the process, from which we obtained the significance of the β coefficient. A significant β means the group status is related to the cell count intensity at the specified location. The z-values in our summary tables correspond to this β coefficient normalized by its sample standard deviation, which under the null hypothesis of no group effect, has an asymptotic standard normal distribution. The p-values give us the probability of obtaining a β coefficient as extreme as the one observed by chance assuming this null hypothesis is true. In the case of three (or more) groups, we utilize Tukey’s Honest Significance test to adjust the p-values of the group factor coefficients to control for multiple comparisons: group1v2, group1v3 and group2v3. To account for multiple comparisons across all voxel/ROI locations, we thresholded the p-values and reported false discovery rates with the Benjamini-Hochberg procedure ^64^. In contrast to correcting for type I error rates, this method controls the number of false positives among the tests that have been deemed significant.

### Stereotaxic injections

All surgery was performed under aseptic conditions and body temperature was maintained with a heating pad. Standard surgical procedures were used for stereotaxic injection and implantation, as previously described^26^. Briefly, mice were anesthetized with isoflurane (2% in a mixture with oxygen, applied at 1.0 L/min), and head-fixed in a stereotaxic injection frame (Stereotax model). Ketamine was used as an anesthetic.

To prepare mice for the photometry experiments, we first made a small craniotomy in each mouse, unilaterally. We then lowered a glass micropipette (tip diameter, ∼20 μm) containing viral solution to reach the ACC (coordinates: +0.55 mm posterior to Bregma, 0.3 mm lateral from midline, and −0.9 mm ventral from brain surface). The injection coordinates for the LC experiments were (+1.5mm posterior to lambda, 0.8mm lateral from the midline, and 2.8mm ventral from the brain surface). (About 0.2–0.3 μL of viral solution was delivered with pressure applications (5–20 psi, 5–20 ms at 1 Hz) controlled by a Picrospritzer and a pulse generator. The rate of injection was ∼50 nl/min. The pipette was left in place for 5-10 minutes following the injection, and then slowly withdrawn. Infection of ACC was performed in both hemispheres in mice dedicated to chemogenetic inhibition experiments and about 0.2 μL of viral solution was delivered to each hemisphere (coordinates: +0.55 mm posterior to Bregma, ± 0.3 mm lateral from midline, and −0.9 mm ventral from brain surface).

We then implanted optic fibers above injection locations in the mice dedicated for photometry experiments (coordinates: +0.55 mm anterior to Bregma, 0.3 mm lateral from midline, and 0.9 mm vertical from brain surface). A head-bar was also mounted for head-restraint. We waited one week for the mice to recover from surgery and pair them with a mate. We then waited for the pups to be born to start recording the photometry signals.

### Fiber photometry recordings and data analysis

To record the activity of CamkII+ cells in ACC *in vivo*, we used a custom-made fiber photometry system to measure GCaMP6s signals in these neurons through an optical fiber (Fiber core diameter, 200 µm; Fiber length, 2.0 mm; NA, 0.37; Inper, Hangzhou, China) unilaterally implanted in ACC of 8-10 weeks-old male and female mice. Animals were habituated to the behavioral box in their home cage for 10 min starting at least 1 day before parturition. 4 retrieval sessions were recorded each day were all pups from the litter were scattered in the cage, and the test subject was allowed to retrieve all the pups. Each session was 5 minutes long and sessions were separated by 2 minutes. For the second pregnancy experiment on supplementary figure 5, we repeated the same procedure with the second litter of pups. The same behavioral procedures were used to record neural activity from all of the populations described below. GCaMP signals were detected and measured as follows: briefly, activity-dependent GCaMP was delivered by AAV to CamkII+ neurons under the expression of the CamkII promoter. An optical fiber cable was mated to the fiber implant in the ACC neurons before each optical recording session, and it was used to deliver 470 nm and 565 nm excitation light to the brain. The intensity of the light for excitation was adjusted to ∼5-10 µW at the tip of the patch cord. The two wavelengths were sinusoidally modulated at 211 Hz and 180 degrees out of phase. Green and red emitted light signals were filtered and split to separate photodetectors and digitally sampled at 6100 Hz via a data acquisition board (National Instruments, Model # NI USB-6211). Peaks were extracted by custom Matlab software with an effective sampling rate of 211 Hz. Each signal was corrected for photobleaching by fitting the decay with a double exponential, and then normalized to a Z score. After subtracting the activity-independent red signal to correct for movement artifacts, the green signal was transformed back to absolute fluorescence and DF/F was computed relative to the mean of the measured fluorescence minus the mean of the baseline fluorescence. The resulting traces from each recording session were converted to a Z score to compare between subjects and across days. All data analysis was performed using custom written code in Matlab.

To record activity of DBH+ cells in LC *in vivo*, we injected a cre-dependent AAV GCaMP7s into the LC of DBH-Cre mice and implanted a fiber unilaterally in the LC (Fiber core diameter, 200 µm; Fiber length, 5.5 mm; NA, 0.37; Inper, Hangzhou, China). The intensity of the light for excitation was adjusted to ∼30 µW at the tip of the patch cord. To record the activity of inhibitory neurons in the ACC during parental behavior, we injected a cre-dependent AAV GCaMP7s into ACC of VGAT-Cre mice and implanted a fiber unilaterally in ACC (Fiber core diameter, 200 µm; Fiber length, 2.0 mm; NA, 0.37; Inper, Hangzhou, China). The intensity of the light for excitation was adjusted to ∼5-10 µW at the tip of the patch cord. To record noradrenaline release in ACC, we used fiber photometry and a noradrenaline sensor. We injected AAV-hsyn-NE2h into the cingulate cortex of CBA/CaJ mice and implanted a fiber unilaterally in the ACC (Fiber core diameter, 200 µm; Fiber length, 2.0 mm; NA, 0.37; Inper, Hangzhou, China). The intensity of the light for excitation was adjusted to ∼5-10 µW at the tip of the patch cord.

To record the activity of LC-ACC neurons *in vivo,* we injected a retrograde AAV.hsyn.CRE.WPRE in ACC of female CBA/CaJ mice, and a cre-dependent AAV GCaMP7s into the LC and implanted a fiber unilaterally in LC (Fiber core diameter, 200 µm; Fiber length, 5.5 mm; NA, 0.37; Inper, Hangzhou, China). The intensity of the light for excitation was adjusted to ∼30 µW at the tip of the patch cord. We recorded 4 retrieval sessions each day. All pups from the litter were scattered in the cage, and the test subject was allowed to retrieve all the pups. Each session was 5 minutes long and sessions were separated by 2 minutes.

### Chemogenetic Inhibition

Mice were habituated to the behavioral box for 10 minutes at least 24h before the experiment. Mice were maintained on a 12h/12 h light-dark cycle (lights on 10:00 h) and received food ad libitum. During the dark phase, test subjects were habituated in their home cage for 10 minutes. CBA/CaJ mice expressing hM4D(Gi) were injected intraperitoneally (i.p.) with either saline (0.9% NaCl) or clozapine (0.1 mg/kg) (HelloBio, Inc.) dissolved in saline. The injection of clozapine or saline was alternating in each mouse every other day; P0 and P2 clozapine and P1 and P3 saline. Twenty minutes after the injections, all pups were scattered in the home cage and the test subject’s behavior was recorded for 10 minutes. Then, all pups were placed in a jar for 5 min and videos were recorded. Lastly, the test subjects were exposed to an empty jar and videos were recoded. All videos were manually scored using BORIS^61^.

For the LC-ACC neuron selective inactivation experiments, we bilaterally injected a retrograde AAV.hsyn.CRE.WPRE in ACC of female CBA/CaJ mice, and cre-dependent inhibitory DREADDS (pAAV.hsyn.DIO.hM4D(Gi).mcherry) in LC. We waited for the mice to recover for a week and paired them with a mate. When the pups were born, we recorded interactions from PND0-PND3. Each day, we habituated the mice to a soundproof box in the home cage with the pups for 10 min. We then injected either saline (0.9% NaCl) or clozapine (0.1 mg/kg) (HelloBio, Inc.) dissolved in saline and waited 20 minutes. Then, we recorded interactions with pups for 10 minutes and quantified the mice latency to retrieve the pups back to the nest.

---

**Supplementary Figure 1.**
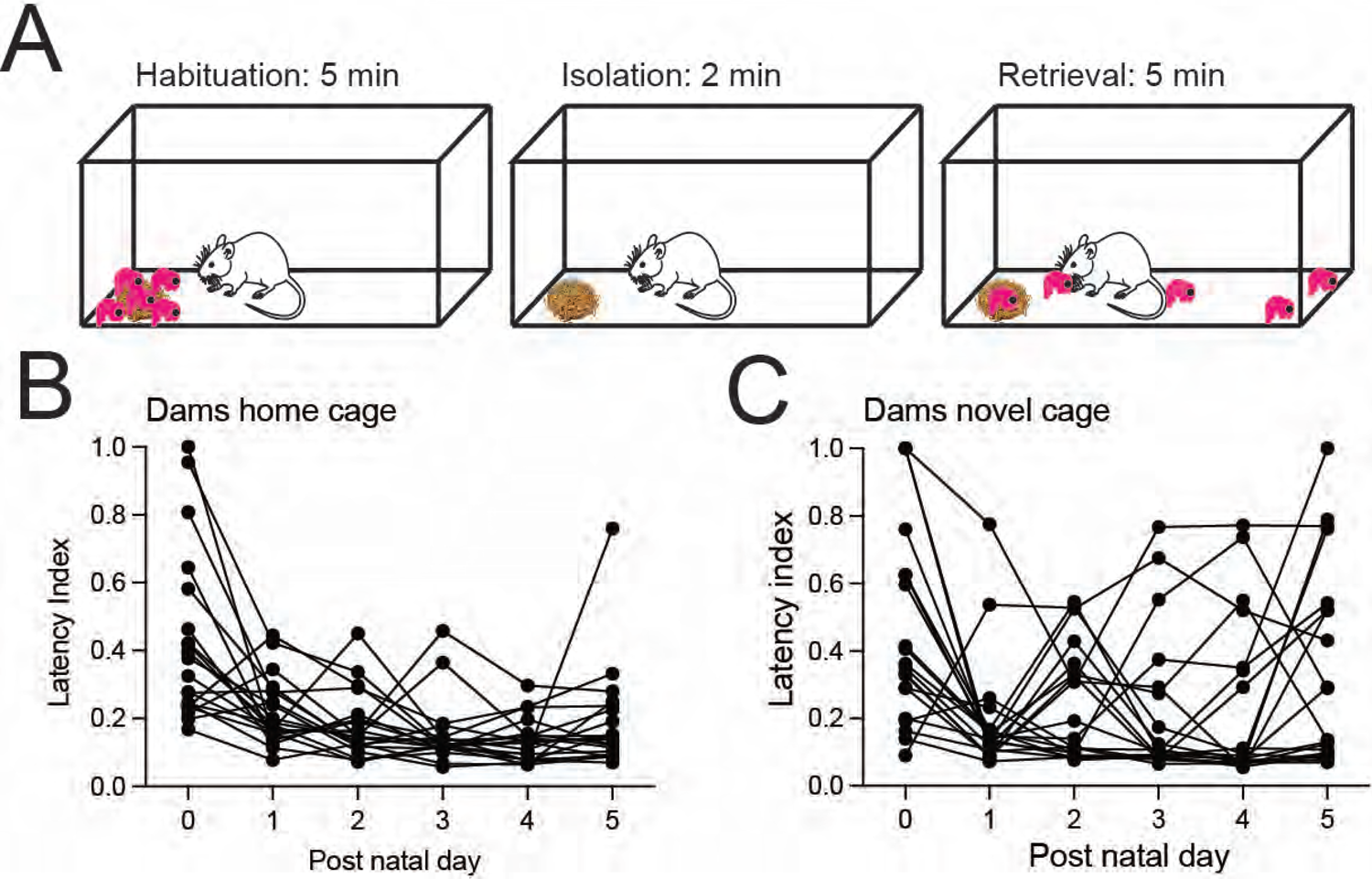
Dams pup retrieval behavior across days. A) Schematic of behavioral paradigm. B) Plot of retrieval latency of dams in the home cage across days. Lines track each individual’s performance across PNDs 0 – 5. C) Plot of retrieval latency of dams in the novel cage across days. Lines track each individual’s performance across PNDs 0 – 5.

**Supplementary Figure 2.**
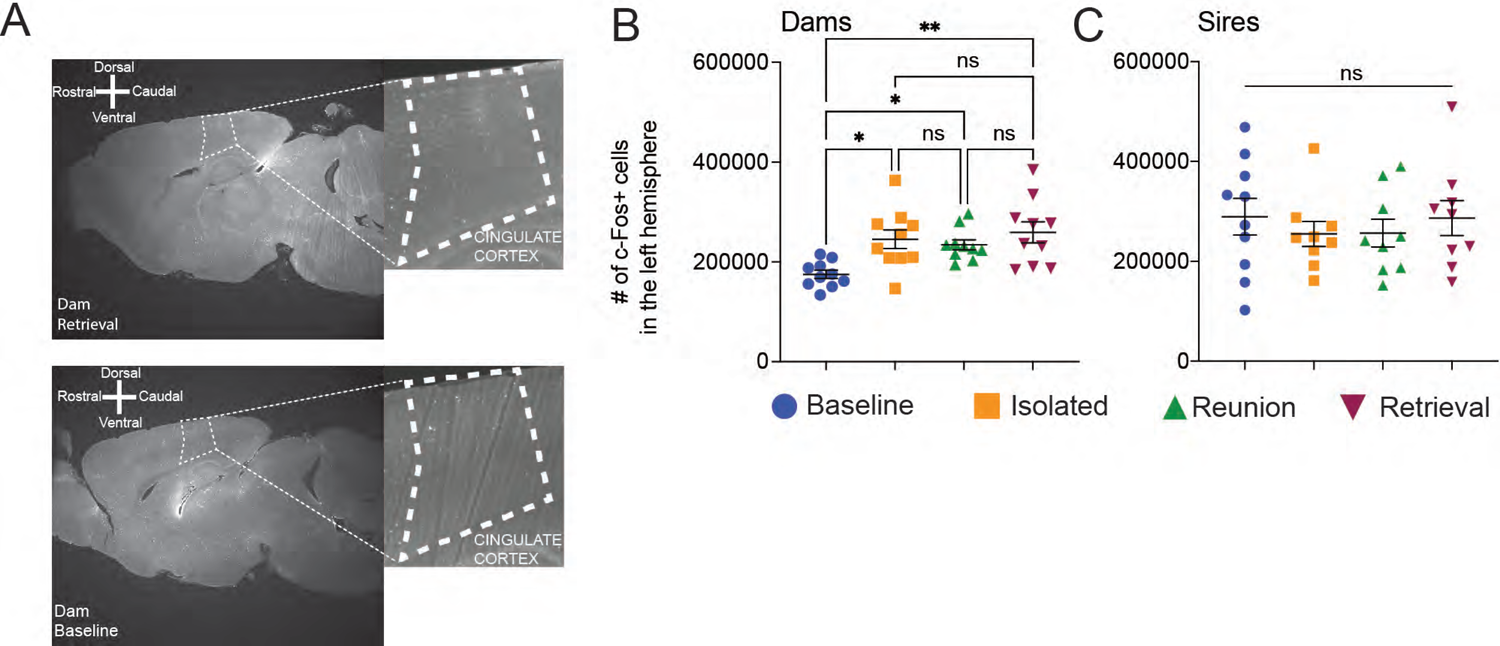
Brain-wide *c-fos*^+^ cell count variability in dams and sires. A) Representative photomicrograph of a sagittal brain optical section showing *c-fos* expression of two dams. The top panel is a dam from the retrieval group, and the bottom panel is a dam from the baseline group. B) Plot of *c-fos^+^* cell counts in the left hemisphere of dams. Ordinary one-way ANOVA; **p=0.0026; F=5.718; Tukey’s multiple comparison test, baseline vs isolated *p=0.0141; baseline vs reunion *p=0.0490; baseline vs retrieval **p=0.0026. C) Plot of *c-fos^+^* cell counts in the left hemisphere of sires. Ordinary one-way ANOVA; not significant.

**Supplementary Figure 3.**
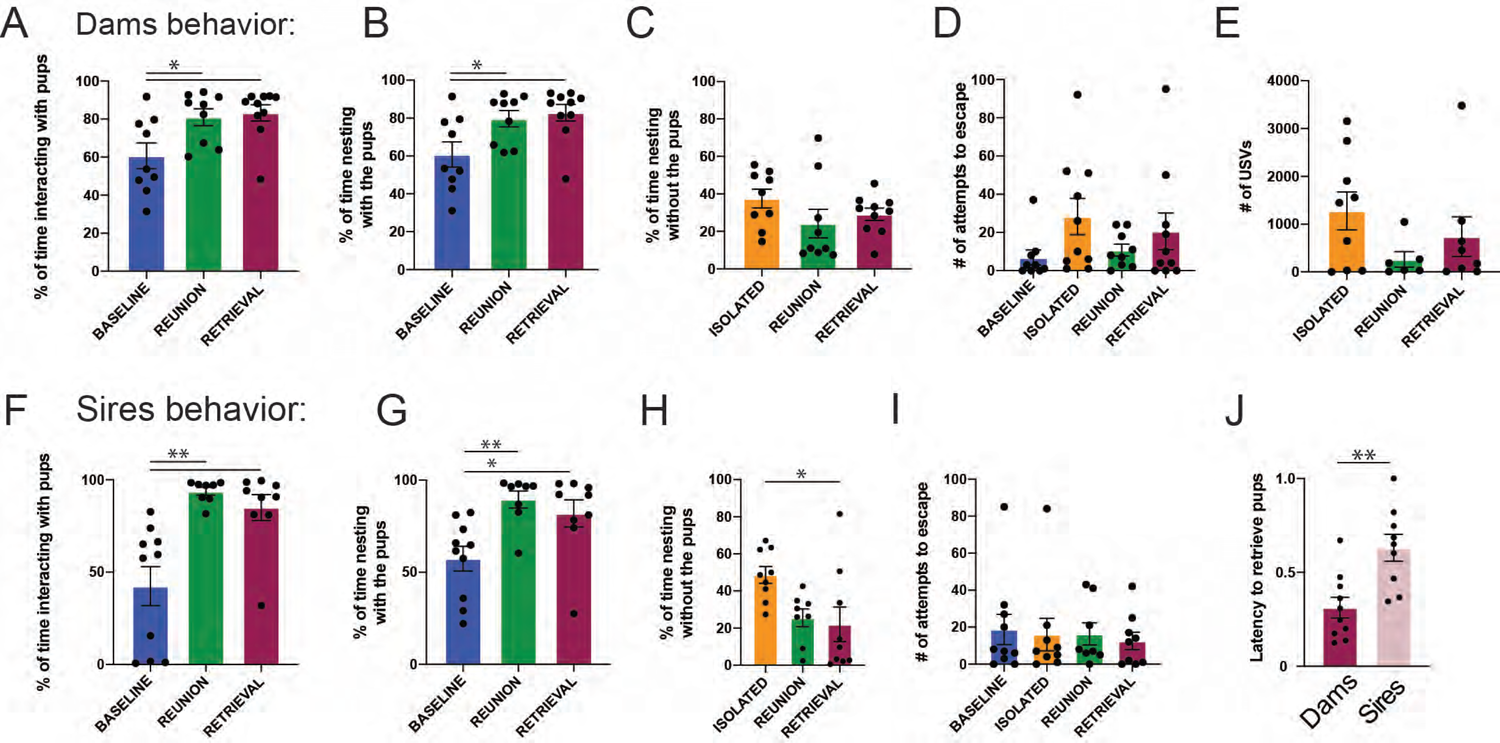
Behavior quantification of dams and sires used for brain-wide *c-fos* mapping. A) Plot of time spent interacting with the pups for the dams’ data set presented as a percentage; lines represent mean ± s.e.m (Ordinary one-way ANOVA **p=0.0098; F=5.596; Tukey’s multiple comparisons test: baseline vs reunion *p=0.0318; baseline vs retrieval *p=0.0133). B) Plot of time spent in the nest with the pups for the dams’ data set presented as a percentage; lines represent mean ± s.e.m (Ordinary one-way ANOVA *p=0.0124; F=5.262; Tukey’s multiple comparisons test: baseline vs reunion *p=0.0455; baseline vs retrieval *p=0.0147). C) Plot of time spent in the nest without the pups for the dams’ data set presented as a percentage; lines represent mean ± s.e.m. (Ordinary one-way ANOVA p=0.2469; F=1.480). D) Plot of total number of escape attempts for the dams’s data set (Ordinary one-way ANOVA p=0.1942; F=1.659). E) Plot of USVs emitted by the dams in the absence of the pups quantified using DeepSqueak; lines represent mean ± s.e.m. (Ordinary one-way ANOVA p=0.1975; F=1.761). F) Plot of time spent interacting with the pups for the sires’ data set presented as a percentage; lines represent mean ± s.e.m (Ordinary one-way ANOVA ***p=0.0002; F=12.19; Tukey’s multiple comparisons test: baseline vs reunion ***p=0.0004; baseline vs retrieval **p=0.0021). G) Plot of time spent in the nest with the pups for the sires’ data set presented as a percentage; lines represent mean ± s.e.m (Ordinary one-way ANOVA ***p=0.0046; F=6.788; Tukey’s multiple comparisons test: baseline vs reunion **p=0.0056; baseline vs retrieval *p=0.0306). H) Plot of time spent in the nest without the pups for the sires’ data set presented as a percentage; lines represent mean ± s.e.m. (Ordinary one-way ANOVA *p=0.0190; F=4.733; Tukey’s multiple comparisons test: isolated vs retrieval *p=0.0237). I) Plot of total number of escape attempts for the sires’ data set (Ordinary one-way ANOVA p=0.9408; F=0.1313). J) Plot showing the latency index of dams and sires of the retrieval group; lines represent mean ± s.e.m (Mann Whitney test **p=0.0041).

**Supplementary Figure 4.**
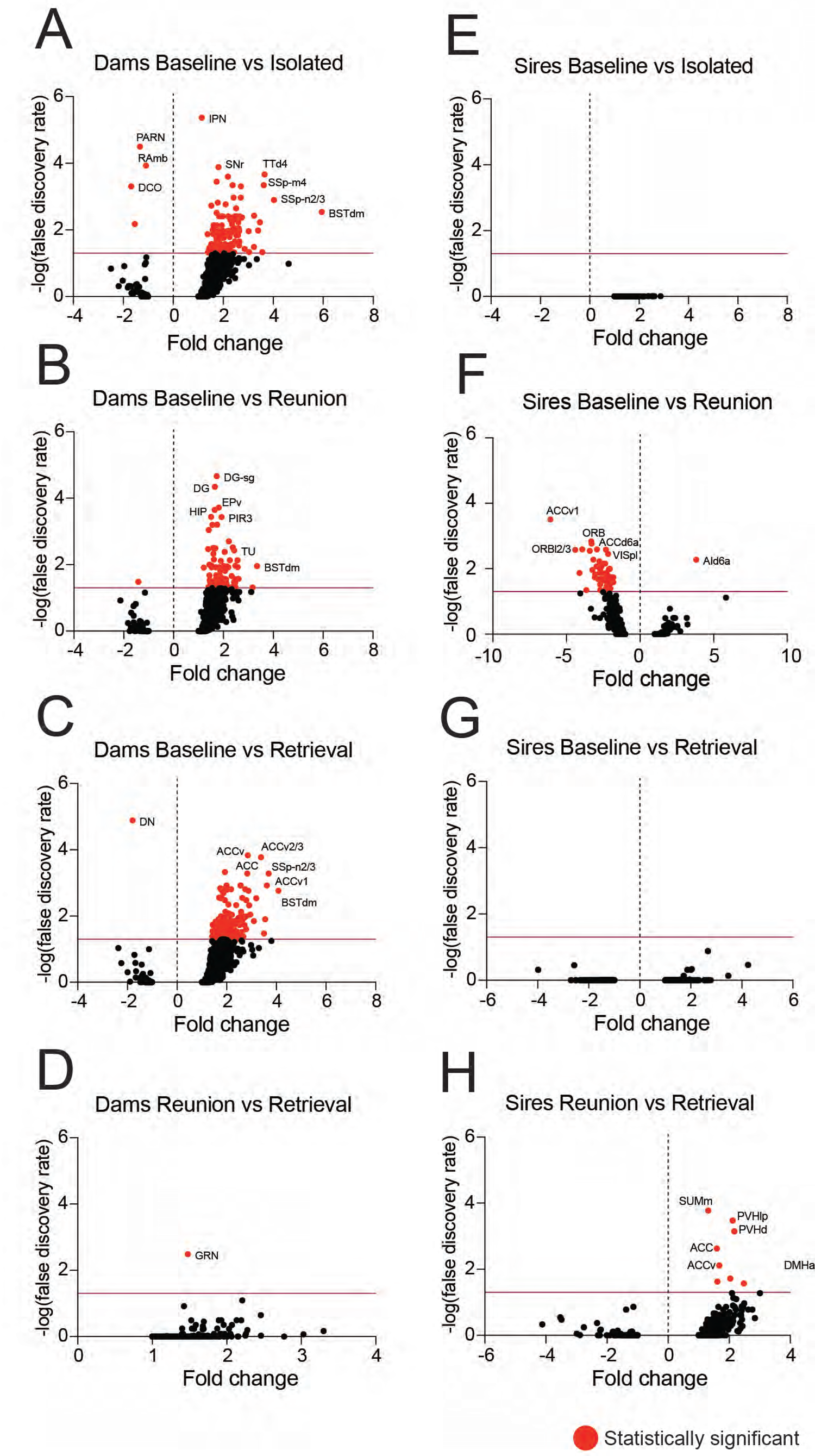
Volcano plots of regions induced by isolated, reunion and retrieval conditions in dams and sires as fold change in *c-fos*+ cells. A) Volcano plot of *c-fos* induction in regions of the baseline vs isolated conditions in dams as fold change in *c-fos*+ cells. B) Volcano plot of *c-fos* induction in regions of the baseline vs reunion conditions in dams as fold change in *c-fos*+ cells. C) Volcano plot of *c-fos* induction in regions of the baseline vs retrieval conditions in dams as fold change in *c-fos*+ cells. D) Volcano plot of *c-fos* induction in regions of the reunion vs retrieval conditions in dams as fold change in *c-fos*+ cells. E) Volcano plot of *c-fos* induction in regions of the baseline vs isolated conditions in sires as fold change in *c-fos*+ cells. F) Volcano plot of *c-fos* induction in regions of the baseline vs reunion conditions in sires as fold change in *c-fos*+ cells. G) Volcano plot of *c-fos* induction in regions of the baseline vs retrieval conditions in sires as fold change in *c-fos*+ cells. H) Volcano plot of *c-fos* induction in regions of the reunion vs retrieval conditions in sires as fold change in *c-fos*+ cells. For all plots, the false discovery rate (FDR) analysis was carried out by the Benjamini-Hochberg procedure. Red data points correspond to statistically significant ROIs. The purple horizontal line indicates the significant threshold with an FDR of 0.05. All ROIs above the line are statistically significant.

**Supplementary Figure 5.**
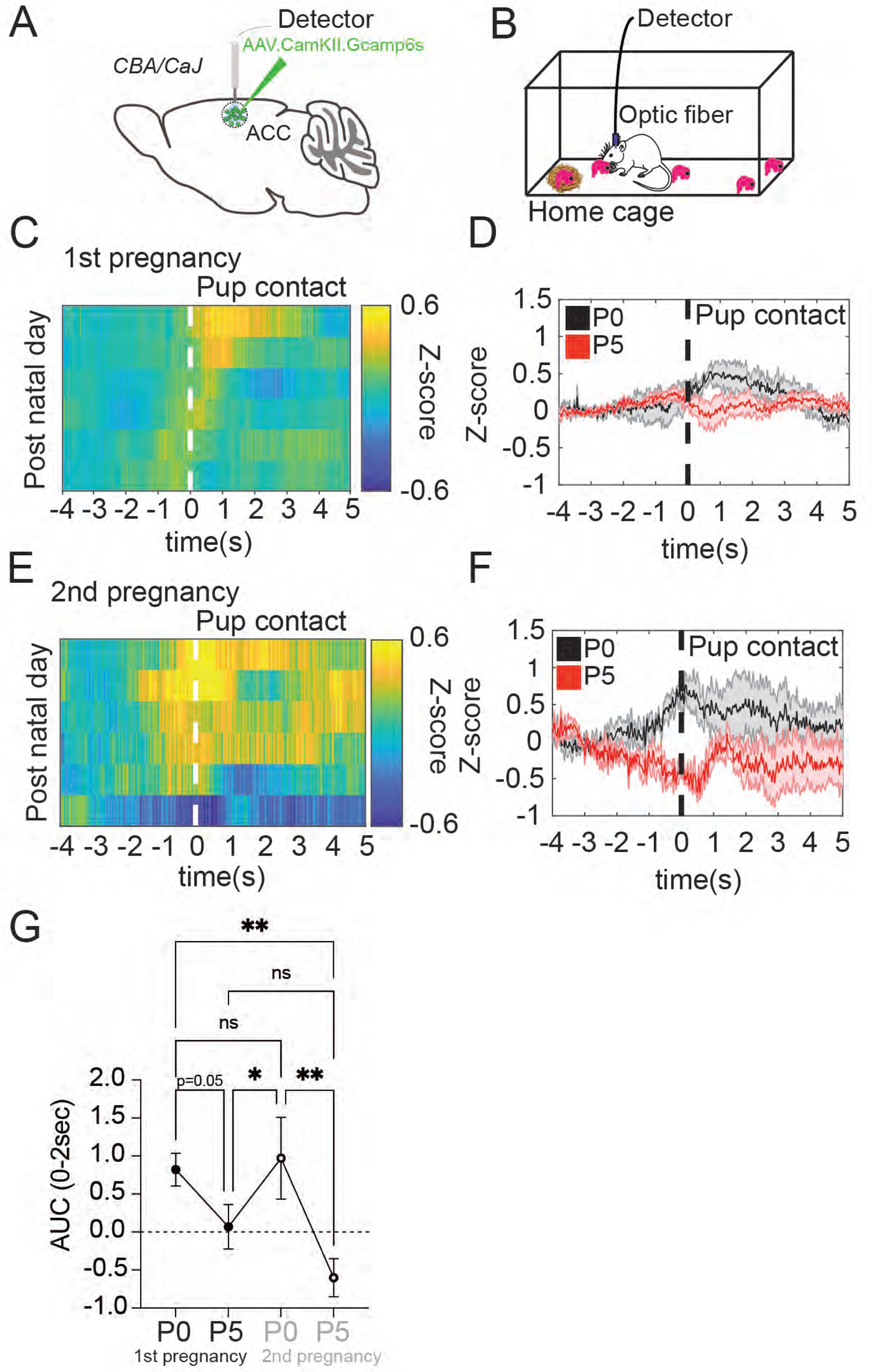
ACC^CAMKII^ neurons calcium transients re-appear on early postnatal days in recordings from a second litter. A) Schematic depicting our viral strategy to express GCaMP6s in excitatory neurons in the anterior cingulate cortex of dams. B) Schematic of behavioral paradigm. C) Heatmap of mean GCaMP6s fiber photometry signals from excitatory neurons in the cingulate cortex during pup gathering events from a first litter (n = 4). The data are aligned to the pup contact. Each row is the mean activity across all mice for each of 6 days. **D)** Plot of the mean Z-scored traces of excitatory neurons for all mice, contrasting post-natal day 0 (black) and post-natal day 5 (red) from a first litter. The panel shows the activity aligned to the pup contact. E) Heatmap of mean GCaMP6s fiber photometry signals from excitatory neurons in the cingulate cortex during pup gathering events from a second litter (n = 4). The data are aligned to the pup contact. Each row is the mean activity across all mice for each of 6 days. **F)** Plot of the mean Z-scored traces of excitatory neurons for all mice, contrasting post-natal day 0 (black) and post-natal day 5 (red) from a second litter. The panel shows the activity aligned to the pup contact. **G)** Comparison of the mean magnitude of the retrieval-related activity of excitatory neurons between PND0 and PND5 from first and second litters, quantified as the mean area under the curve from pup contact to 2 seconds after of traces showing a decline in the magnitude of activity over post-natal days 0 – 5. Repeated measures ANOVA (**p=0.0039; F=9.409; Benjamini, Krieger and Yekutieli multiple comparison test, PND0-first-pregnancy vs PND5-first-pregnancy p=0.0516; PND0-1st-pregnancy vs. PND5-2nd-pregnancy **P=0.0022; PND5-1st-pregnancy vs. PND0-2nd-pregnancy *p=0.0247; PND0-2nd-pregnancy vs. PND5-2nd-pregnancy **p=0.0011).

**Supplementary Figure 6.**
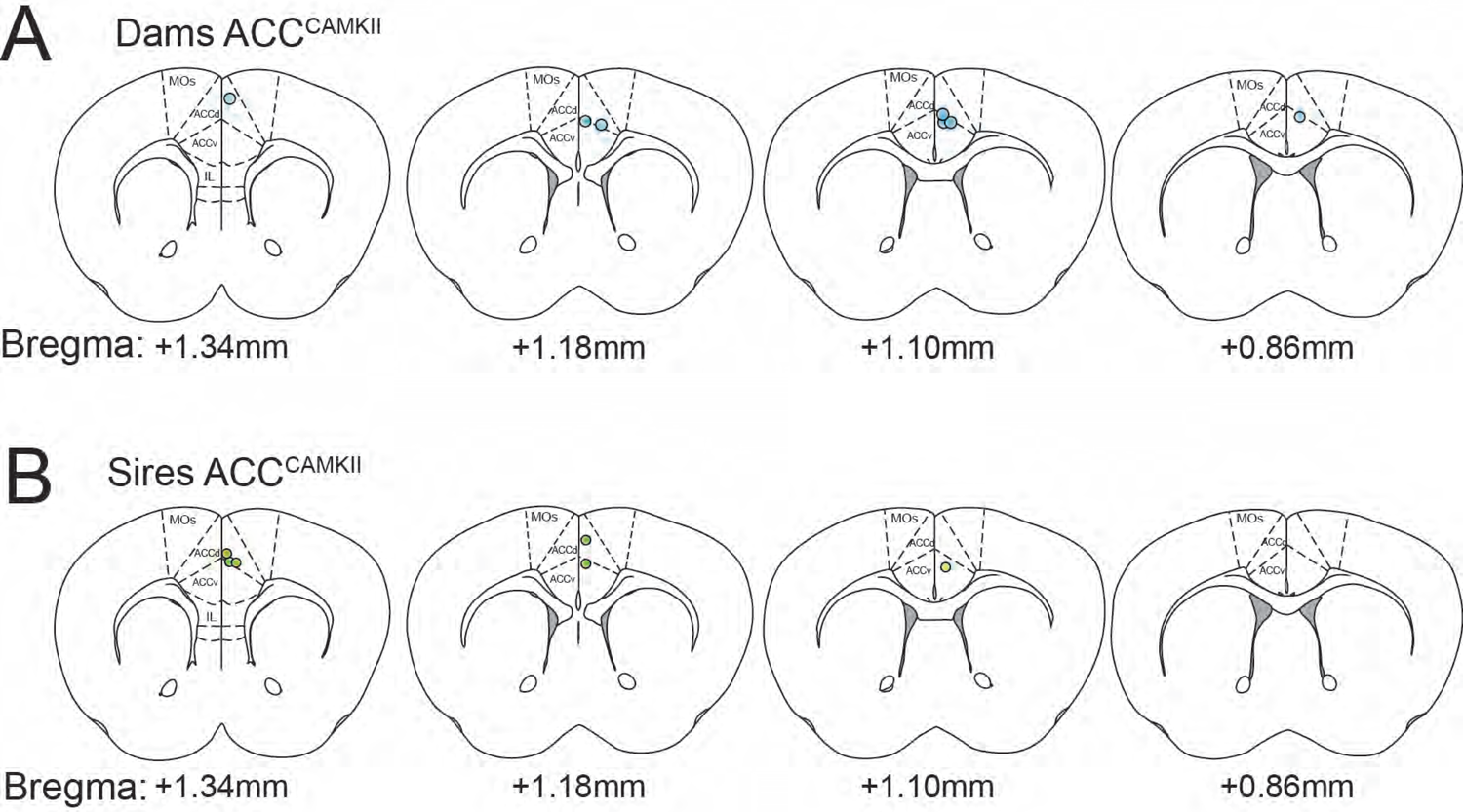
Fiber placements for ACC^CAMKII^ neural recordings in dams and sires. A) Schematic of optic fiber placement in dams ACC^CAMKII^ neurons used in figures 3 and 4 (n=7). B) Schematic of optic fiber placement in sires ACC^CAMKII^ neurons used in figures 3 and 4 (n=6).

**Supplementary Figure 7.**
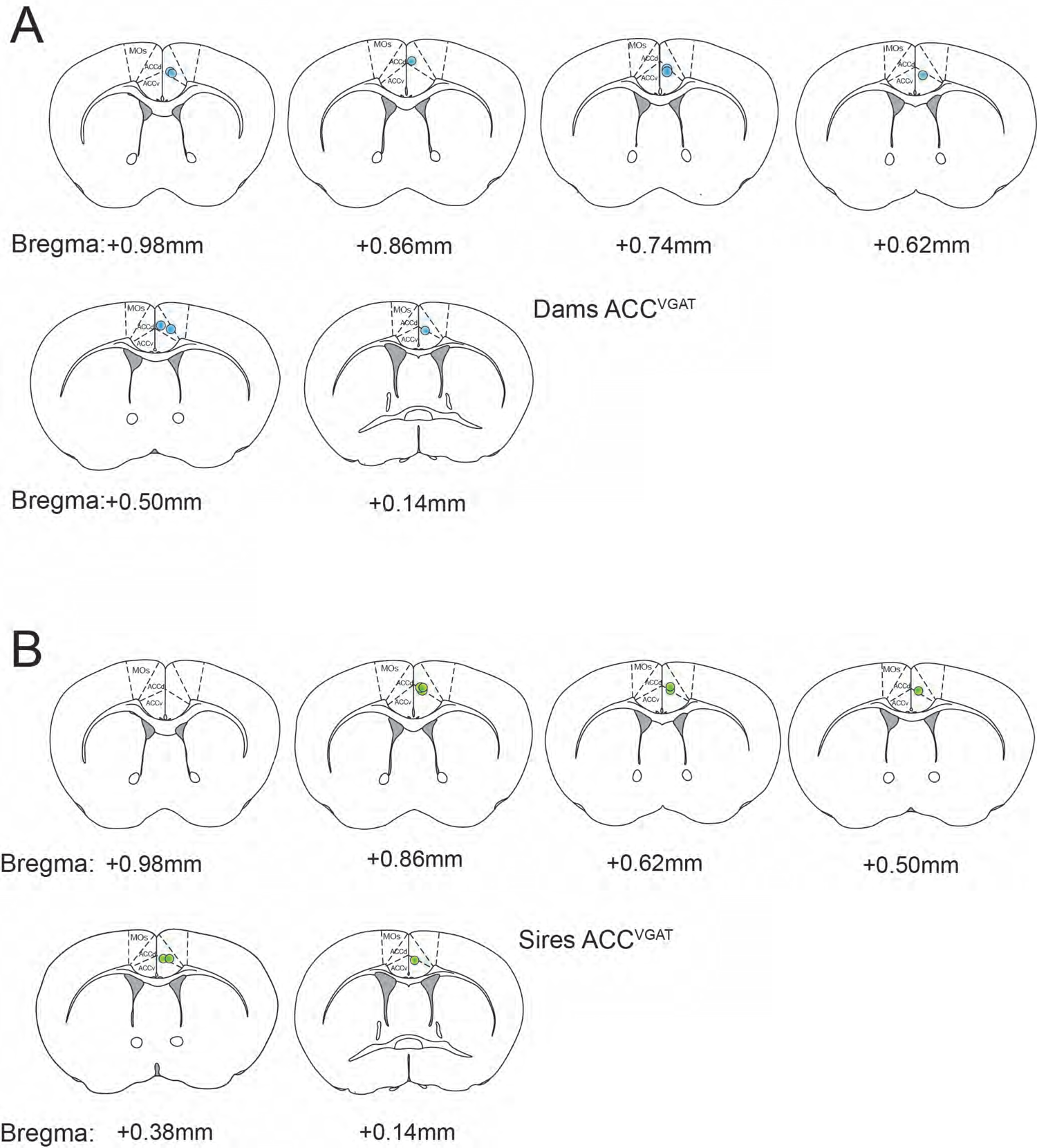
Fiber placements for ACC^VGAT^ neural recordings in dams and sires. A) Schematic of optic fiber placement in dams ACC^VGAT^ neurons used in figures 3 and 4 (n=9). B) Schematic of optic fiber placement in sires ACC^VGAT^ neurons used in figures 3 and 4 (n=9).

**Supplementary Figure 8.**
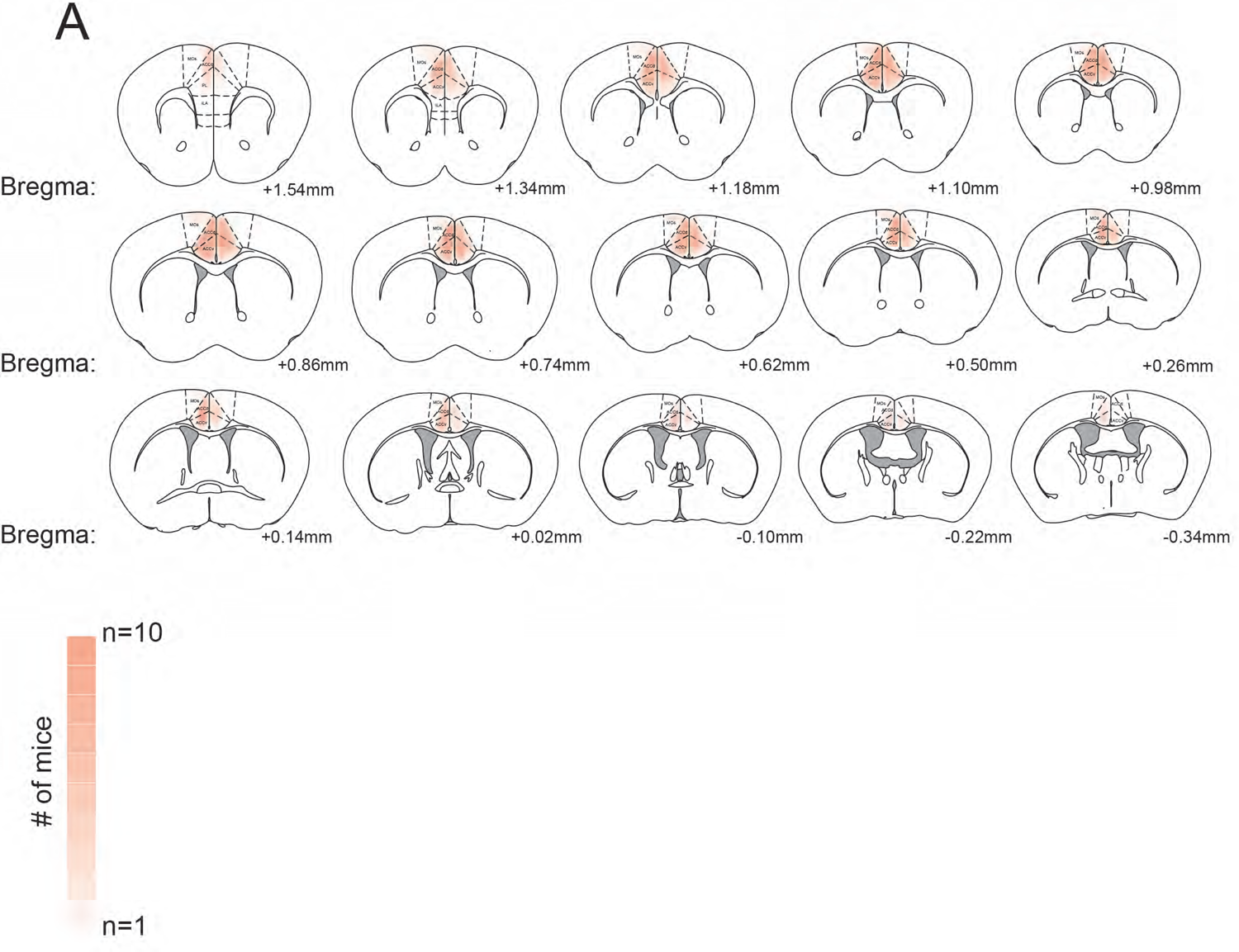
Viral expression for ACC^CAMKII^ chemogenetic inhibition experiment in dams. A) Schematic of viral expression in dams ACC^CAMKII^ neurons used in figure 5 (n=12). The intensity of pink labeling represents overlapping expression in multiple mice.

**Supplementary Figure 9.**
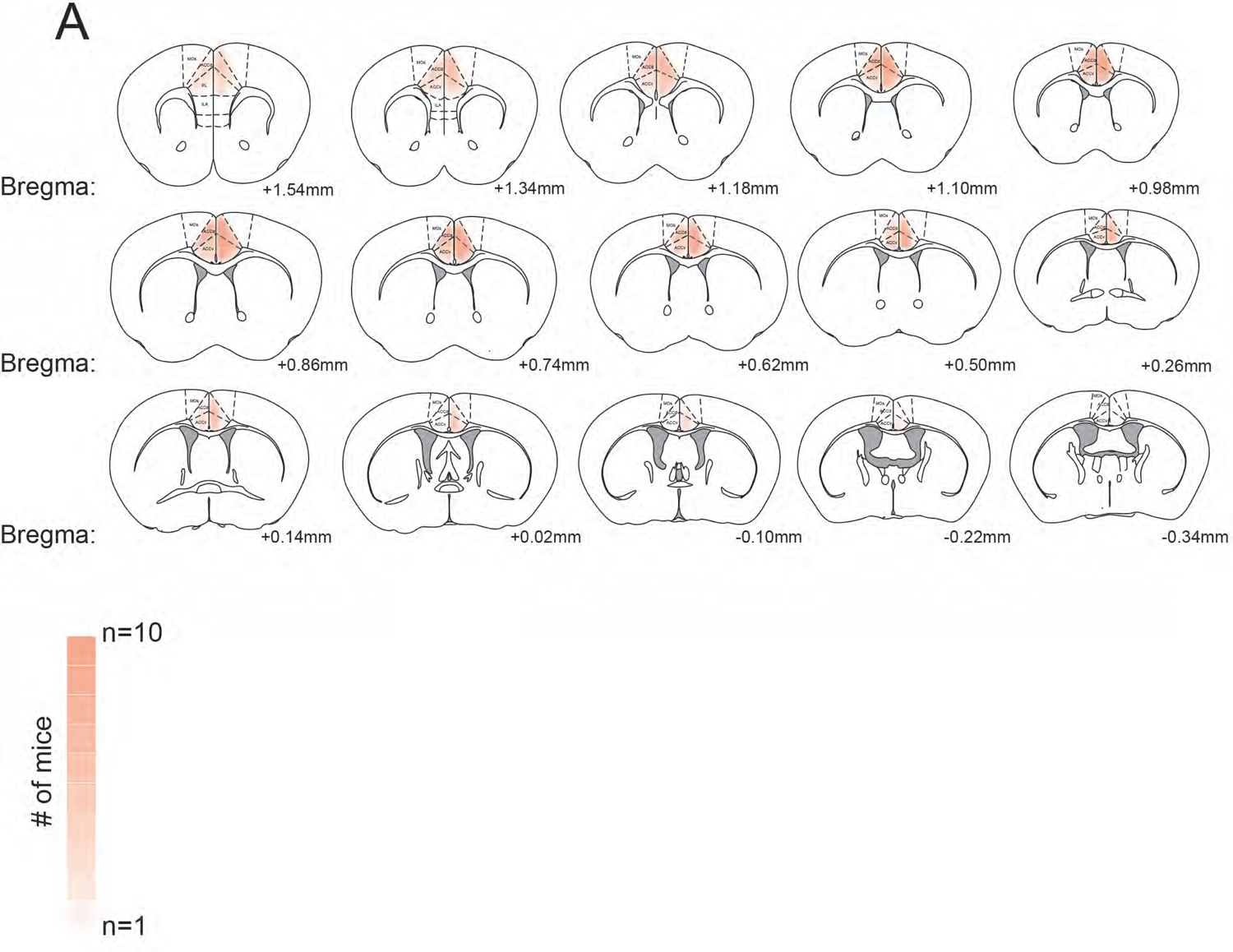
Viral expression for ACC^CAMKII^ chemogenetic inhibition experiment in sires. A) Schematic of viral expression in sires ACC^CAMKII^ neurons used in figure 5 (n=9). The intensity of pink labeling represents overlapping expression in multiple mice.

**Supplementary Figure 10.**
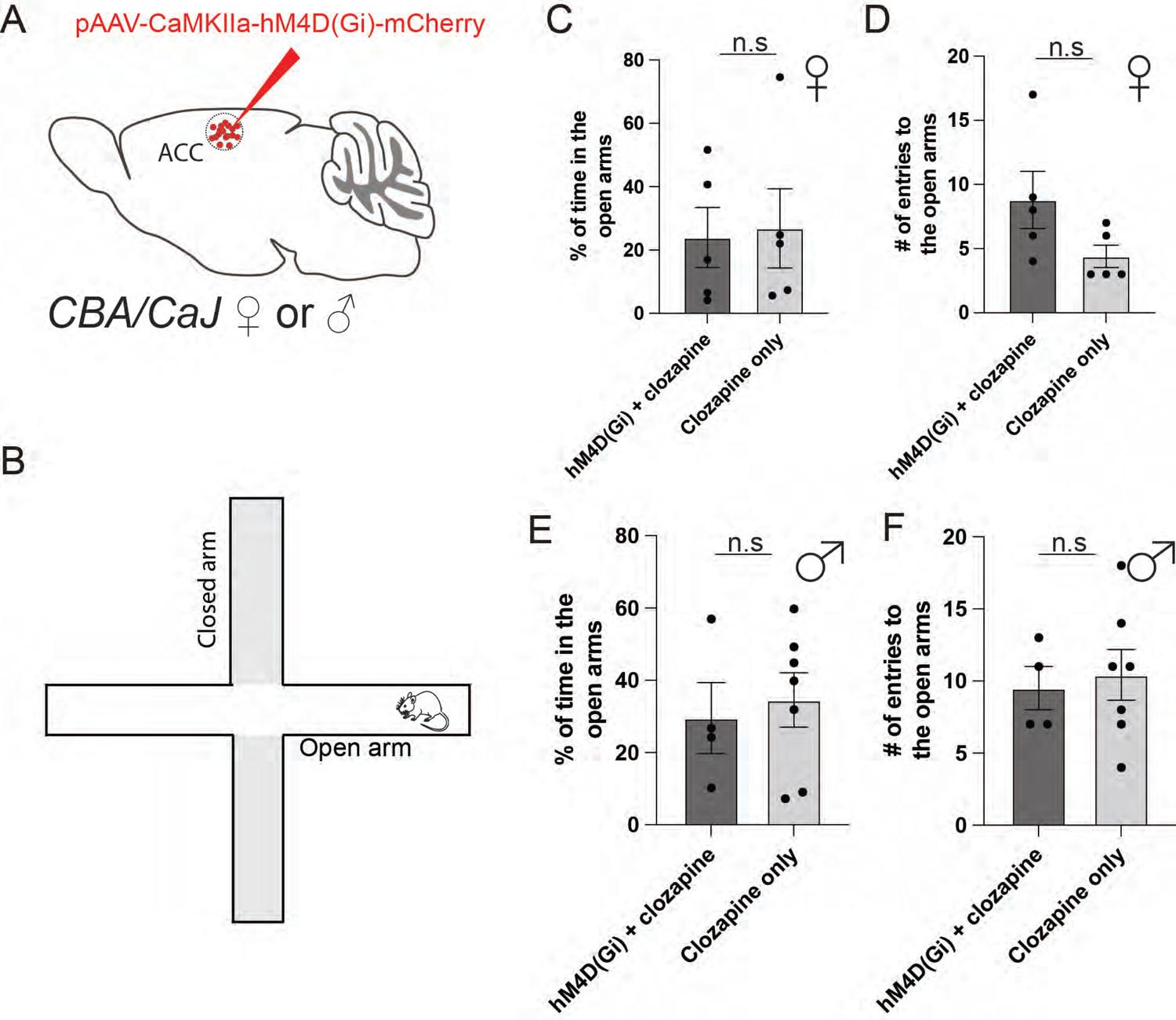
Inhibition of excitatory neurons in the ACC does not alter anxiety-like behaviors in the elevated plus maze. A) Viral strategy to inhibit the ACC excitatory cell population using chemogenetics. B) Schematic of the elevated plus maze setup. C) Plot of the percent of time spent in the open arms by the dams (hM4D(Gi) n=5; control n=5; unpaired t-test not significant p=0.8591). D) Plot of the number of entries to the open arms by the dams (hM4D(Gi) n=5; control n=5; unpaired t-test not significant p=0.1026). E) Plot of the percent of time spent in the open arms by the sires (hM4D(Gi) n=4; control n=7; unpaired t-test not significant p=0.6981). F) Plot of the number of entries to the open arms by the sires (hM4D(Gi) n=4; control n=7; unpaired t-test not significant p=0.7307).

**Supplementary Figure 11.**
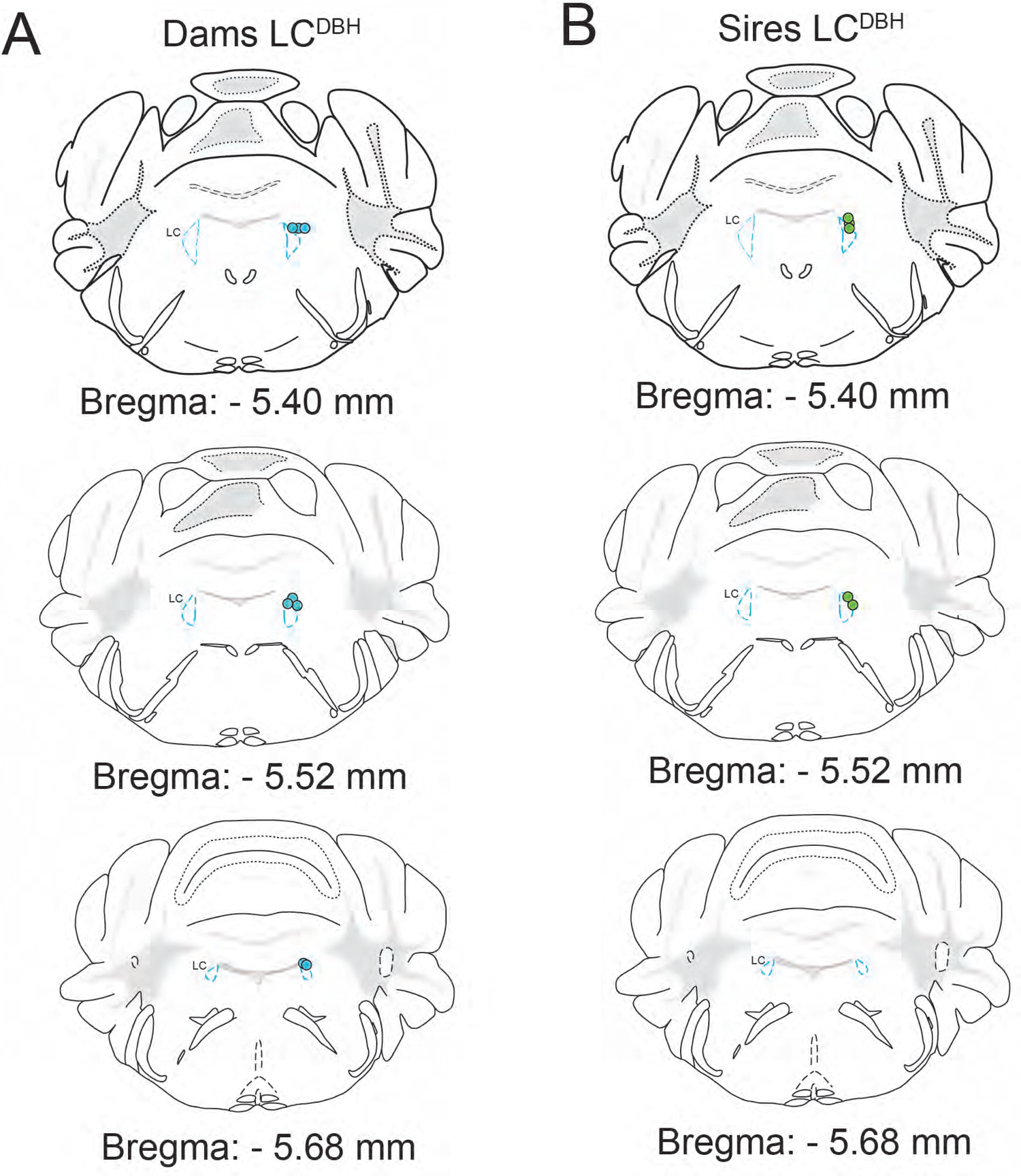
Fiber placements for LC^DBH^ neural recordings in dams and sires. A) Schematic of optic fiber placement in dams LC^DBH^ neurons used in figure 6 (n=8). B) Schematic of optic fiber placement in sires LC^DBH^ neurons used in figure 6 (n=5).

**Supplementary Figure 12.**
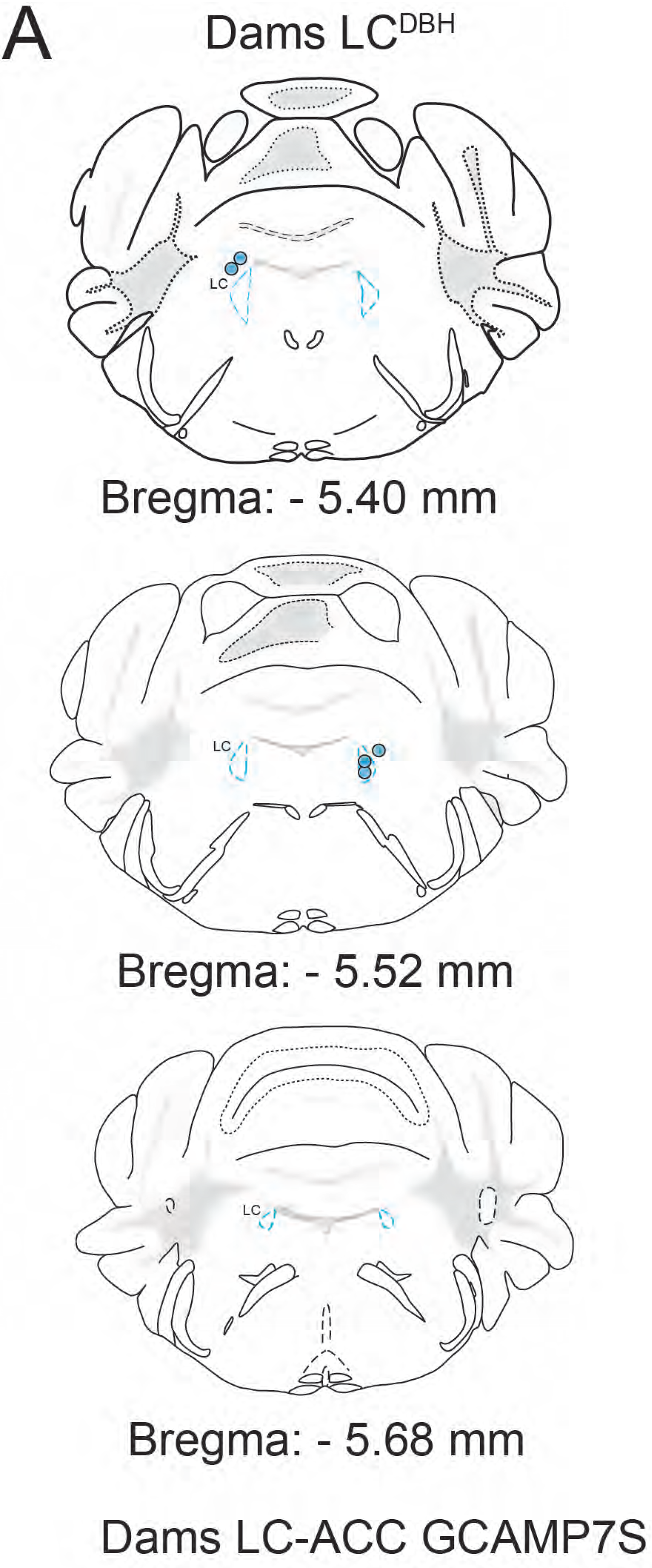
Fiber placements for LC-ACC neural recordings in dams. A) Schematic of optic fiber placement in dams LC-ACC neurons used in figure 7 (n=5).

**Supplementary Figure 13.**
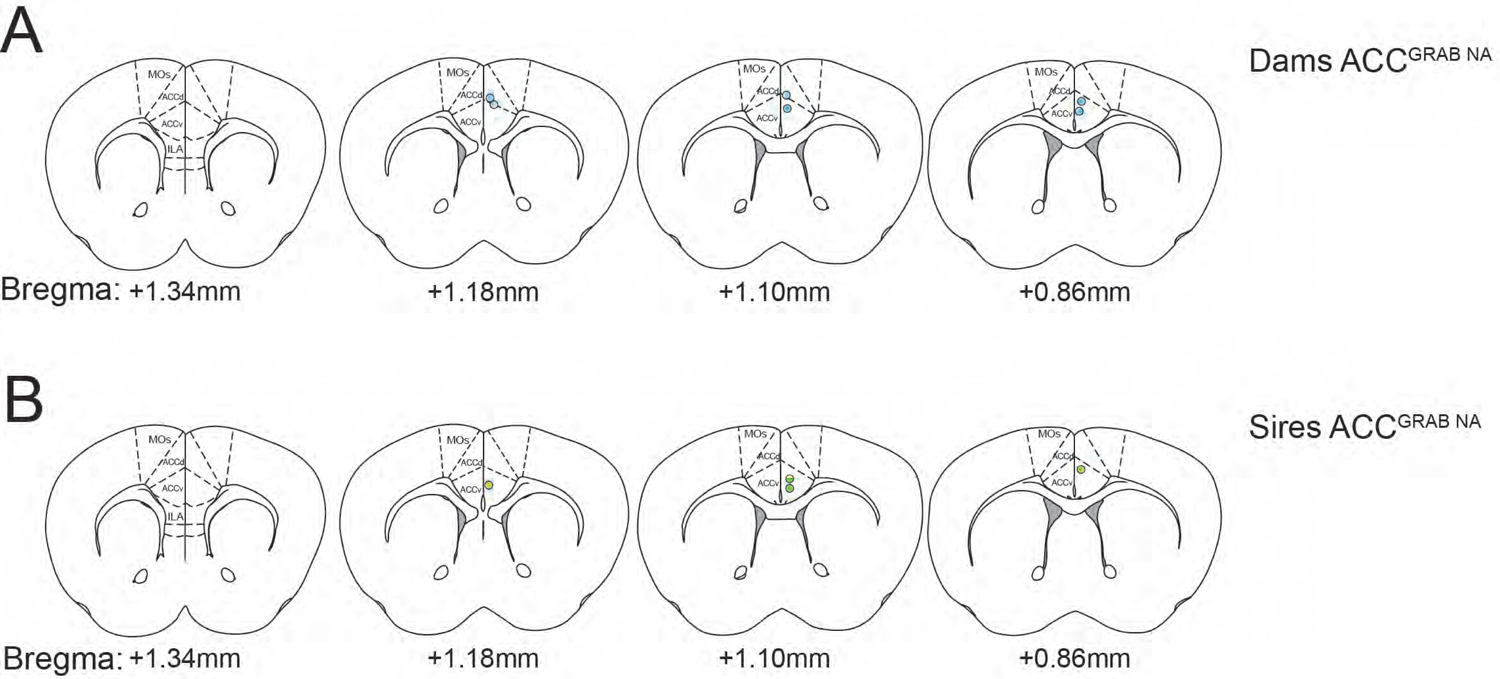
Fiber placements for GRAB^NE^ neural recordings in dams and sires. A) Schematic of optic fiber placement in dams’ GRAB^NE^ recordings used in figure 7 (n=6). B) Schematic of optic fiber placement in sires’ GRAB^NE^ recordings in used in figure 7 (n=4).

